# Shared genetic architecture underlying root metaxylem phenotypes under drought stress in cereals

**DOI:** 10.1101/2020.11.02.365247

**Authors:** Stephanie P. Klein, Jenna E. Reeger, Shawn M. Kaeppler, Kathleen M. Brown, Jonathan P. Lynch

## Abstract

Root metaxylem are phenotypically diverse structures whose function is related to their anatomy, particularly under drought stress. Much research has dissected the genetic machinery underlying metaxylem phenotypes in dicots, but monocots are relatively unexplored. In maize (*Zea mays*), a robust pipeline integrated a GWAS of root metaxylem phenes under well-watered and water stress conditions with a gene co-expression network to identify candidate genes most likely to impact metaxylem phenotypes. We identified several promising candidate genes in 14 gene co-expression modules inferred to be functionally relevant to xylem development. We also identified five gene candidates that co-localized in multiple root metaxylem phenes in both well-watered and water stress conditions. Using a rice GWAS conducted in parallel, we detected overlapping genetic architecture influencing root metaxylem phenotypes by identifying eight pairs of syntenic candidate genes significantly associated with metaxylem phenes. There is evidence that the genes of these syntenic pairs may be involved in biosynthetic processes related to the cell wall, hormone signaling, oxidative stress responses, and drought responses. Our study demonstrates a powerful new strategy for identifying promising gene candidates and suggests several gene candidates that may enhance our understanding of vascular development and responses to drought in cereals.

**One sentence summary:** Cross-species genome-wide association studies and a gene coexpression network identified genes associated with root metaxylem phenotypes in maize under water stress and non-stress and rice.

## Introduction

Drought is a primary constraint to crop production worldwide and its effects are only predicted to worsen in the coming decades as a result of climate change (Lobell et al., 2011; Dai, 2013; IPCC, 2013). Plant breeding efforts have been aimed at maintaining crop productivity in stressful environments in order to sustain growing agriculture-centric economies and to meet increasing consumer demand. As the primary organ responsible for the sensing and uptake of water, roots have become a focus for crop improvement under drought (Lynch et al., 2014; Vadez, 2014). Recent efforts have sought to approach root-focused breeding by selecting for a root system architecture better adapted to water-limited conditions (Ho et al., 2005; Lynch, 2013; Li et al., 2018; Zhang et al., 2019; Burridge et al., 2020). For instance, a steep root growth angle that increases rooting depth has been associated with improved tolerance to water scarcity in maize (Liakat Ali et al., 2015; Pires et al., 2020) and rice (Uga et al., 2013). Fewer nodal roots with more sparsely distributed long lateral roots is also associated with improved drought tolerance (Zhan et al., 2015; Gao and Lynch, 2016; Lynch, 2019). Several anatomical phenes have been proposed to mitigate drought effects by facilitating the capture of mobile resources, such as water and nitrogen, deep in the soil profile (Lynch, 2013; Lynch et al., 2014; Lynch, 2018; Lynch, 2019). Increased aerenchyma production (Zhu et al., 2010; Jaramillo et al., 2013) and larger cortical cells arranged in fewer files cheapen the root maintenance costs (Chimungu et al., 2014a; Chimungu et al., 2014b), which would allow more resources to be reallocated to deeper root construction (Lynch, 2013; Lynch et al., 2014). Additionally, many architectural and anatomical phenes are plastic, which may be an adaptive response to drought stress (Kano et al., 2011; Klein et al., 2020; Schneider et al., 2020a; Schneider and Lynch, 2020; Schneider et al., 2020b). However, less attention has been paid to root hydraulics, which have direct implications for water uptake and transport (Wasson et al., 2012; Vadez, 2014; Maurel and Nacry, 2020).

Maize, one of the world’s most economically and nutritionally important crops, is particularly sensitive to drought stress (Lobell et al., 2014; Daryanto et al., 2016). Root metaxylem vessels are a natural target to improve maize since their size and number in the root is related to abiotic stress tolerance (Richards and Passioura, 1989; Comas et al., 2013). Several studies have found an association between drought tolerance and metaxylem vessel number (de Souza et al., 2013; Klein et al., 2020) or metaxylem vessel area (Richards and Passioura, 1989; Abd Allah et al., 2010; Purushothaman et al., 2013; de Souza et al., 2013; Klein et al., 2020). Plants with narrower root metaxylem have reduced hydraulic conductivity but more conservative water usage and greater resistance to cavitation (Richards and Passioura, 1989; Sperry and Saliendra, 1994; Vilagrosa et al., 2012; Guet et al., 2015). Given that hydraulic conductance is proportional to the vessel radius raised to the fourth power according to the Hagen-Poiseuille equation, even a slight narrowing of the metaxylem vessels will result in a considerable reduction in axial conductivity. Conservative water usage as a result of restricted hydraulic conductance, especially early in the growing season, has been suggested to be more effective than deeper rooting by making water access more likely during the critical anthesis-silking interval (Zaman-Allah et al., 2011; Feng et al., 2016). Reduced transpiration due to restricted hydraulic conductance from narrow metaxylem vessels may also be beneficial for mitigating drought effects by inducing stomatal closure, increasing water use efficiency by reducing the overall shoot size, and retaining moisture at the root tips to enable further growth through the soil profile (Vadez et al., 2014; Lynch, 2019). A constitutive reduction in metaxylem vessel area could limit plant maximum potential growth or reduce relative growth rates () under well-watered conditions (Wahl and Ryser, 2000; Comas et al., 2013), but these effects may be counteracted by augmenting the number of metaxylem vessels (Lynch et al., 2014). Metaxylem phenes that improve performance under water stress with no penalty under optimal conditions are of utmost importance to identify.

Metaxylem vessels form through the careful coordination of phytohormone biosynthesis and signaling, cell wall formation and lignification, cell expansion, radial patterning, and programmed cell death (Roberts and McCann, 2000; Schuetz et al., 2012; Milhinhos and Miguel, 2013; Růžička et al., 2015). Decades of research have deepened our understanding of the genetic and developmental underpinnings of xylem development but has mostly been limited to dicotyledonous or woody species. Recent studies have identified two quantitative trait loci intervals associated with xylem area traits in spring barley (*Hordeum vulgare*) (Oyiga et al., 2020) and a few genetic loci associated with root metaxylem size and number in rice (*Oryza sativa*) (Kadam et al., 2017). However, the genetic mechanisms regulating xylem development in grain crops is still poorly understood. Arranged in a polyarchy pattern (Hochholdinger, 2009), the vascular organization of monocot roots, such as members of the *Poaceae,* is structurally distinct from di-, tri-, or tetrarch patterns typical of most dicots. Cell wall composition also differs between dicots and monocots, since monocots are richer in hydroxycinnamates and hemicellulose with little pectin or structural protein content (Carpita, 1996; Smith and Harris, 1999; Vogel, 2008). Despite these differences, shared genetic machinery underlying xylem development is likely, as is the case for example for the cellulose synthase A (CesA) gene family, which is present in all seed plants (Richmond and Somerville, 2000). Furthermore, comparisons between maize and closely related species have found shared genetic architecture influencing sorghum (*Sorghum bicolor*) root architecture (Zheng et al., 2020) and rice (*Oryza sativa*) grain development (Chen et al., 2016). These studies suggest there is a high likelihood that the genetic machinery regulating other developmental processes, like xylem development, may be conserved within monocots.

Plasticity in xylem phenotypes in response to drought has been observed in maize (Klein et al., 2020; Schneider et al., 2020a), rice (Henry et al., 2012; Kadam et al., 2017), and wheat (*Triticum aestivum*) (Kadam et al., 2015), and are likely mediated by stress-responsive cellular and genetic factors. The genetic mechanisms underlying drought-induced variation in root xylem phenotypes are largely unknown. Several genes regulating biosynthesis and signaling of brassinosteroid, cytokinin, ethylene, and gibberellin have been directly implicated in constitutive and abiotic stress-responsive xylem development in dicot roots (Milhinhos and Miguel, 2013; Ramachandran et al., 2020). However, these complex signaling cascades facilitated by hormones are likely modified by crosstalk with other phytohormones like abscisic acid (ABA), whose production increases in response to drought (Finkelstein, 2013; Ramachandran et al., 2020). In maize, increased ABA production induced by drought coincides with increased root hydraulic conductivity, which is likely mediated by augmented activity of root aquaporins rather than changes to root metaxylem anatomy (Hose et al., 2000; Parent et al., 2009). More recent work showed that ABA interfered with signaling among key developmental regulators and subsequently altered root xylem phenotypes in *Arabidopsis* under limited water availability (Ramachandran et al., 2018). However, there are likely many more genetic components participating in these signaling cascades that contribute to variation in xylem phenotypes and water uptake.

Novel breeding and phenotyping technologies have increased the capacity to evaluate and map desirable characteristics to genetic loci in large populations, as is done in a genome-wide association study (GWAS). Even so, many challenges remain when analyzing GWAS outputs to avoid false positives and isolate the true loci with significant causal effects on phenotype. One method to increase robustness of an associative mapping study is by conducting a comparative multispecies GWAS, which has the potential to enhance our understanding of shared genetic architectures among species (Chen et al., 2016; Zheng et al., 2020). Although maize and rice are different in many respects (e.g. photosynthesis processes, tillering, etc.), their genomes share a high degree of synteny (Bennetzen and Freeling, 1997; Salse et al., 2004; Buell et al., 2005), and in the particular case of xylem phenes, both species have polyarch vascular arrangements. Performing such a comparative GWAS between two grass species of high synteny, like maize and rice, may reveal genetic architectures conserved among the cereals.

A second method to increase the robustness of GWAS is coupling it with a gene co-expression network, which is a helpful tool to infer a candidate gene’s biological function relative to the functional role filled by comparably expressed genes (Schaefer et al., 2017; Schaefer et al., 2018). Complex quantitative traits are often pleiotropic and are associated with many significant SNPs with small effect sizes (Ingvarsson and Street, 2011). A candidate gene’s topological characteristics within the gene network may provide insight on its biological role and essentiality. For instance, hub genes (genes that interact with a large number of other members of the subnetwork) and bottleneck genes (genes with a high level of betweenness centrality) are posited to be the most essential for proper function (Jeong et al., 2001; Yu et al., 2004; Yu et al., 2007). Modifying or removing the activity of hub and bottleneck genes has the greatest chance of inducing measurable downstream effects. Thus, integrating a GWAS with a gene co-expression network is a powerful method to identify candidate genes with the highest likelihood of shaping a measurable effect on phenotype and function.

Root metaxylem phenotypes merit greater attention because of their potential to improve crop performance under water-limited conditions. One course of action is to dissect their genetic architecture to provide insight on the loci and overarching mechanisms contributing to root metaxylem phenotypic variation under abiotic stress. In this study, we phenotyped the root metaxylem of a large diversity panel of field-grown maize under well-watered and water stress conditions. A parallel study phenotyped the root metaxylem of a rice diversity panel grown in the greenhouse under well-watered conditions (Vejchasarn 2014). Our objectives were to test the hypotheses that 1.) natural variation in root metaxylem phenotypes is under genetic control; 2.) genetic loci associated with root metaxylem phenotypes under WS are unique from those activated under WW; and 3.) there is shared genetic architecture associated with root metaxylem phenotypes in maize and rice.

## Results

### Natural variation in root metaxylem phenotypes in the field

Wide variation in all root metaxylem phenes was observed under both well-watered and water deficit stress (hereafter, water stress) conditions in 180 and 412 genotypes of the Wisconsin Diversity Panel in 2015 and 2016, respectively (Figure 1). A 3-4-fold range in values were observed for all metaxylem phenes under both water treatments except for volumetric flow rate, which exhibited a 31-fold range. Watering regime had no significant overall effect on metaxylem phenotypes except for volumetric flow rate (p = 0.036) (Table 1), but responses to drought varied by genotype (Supplemental Figure S1). Plasticity in all metaxylem vessel phenes in response to water stress ranged from −84%-351% relative to the phene state in well-watered conditions. While total metaxylem vessel area was stable between seasons, there were significant seasonal effects for maximum and median metaxylem vessel areas, metaxylem vessel number, and volumetric flow rate (Table 1). Volumetric flow rate was the only phene to exhibit a significant interactive year and treatment effect.

**Figure 1.**
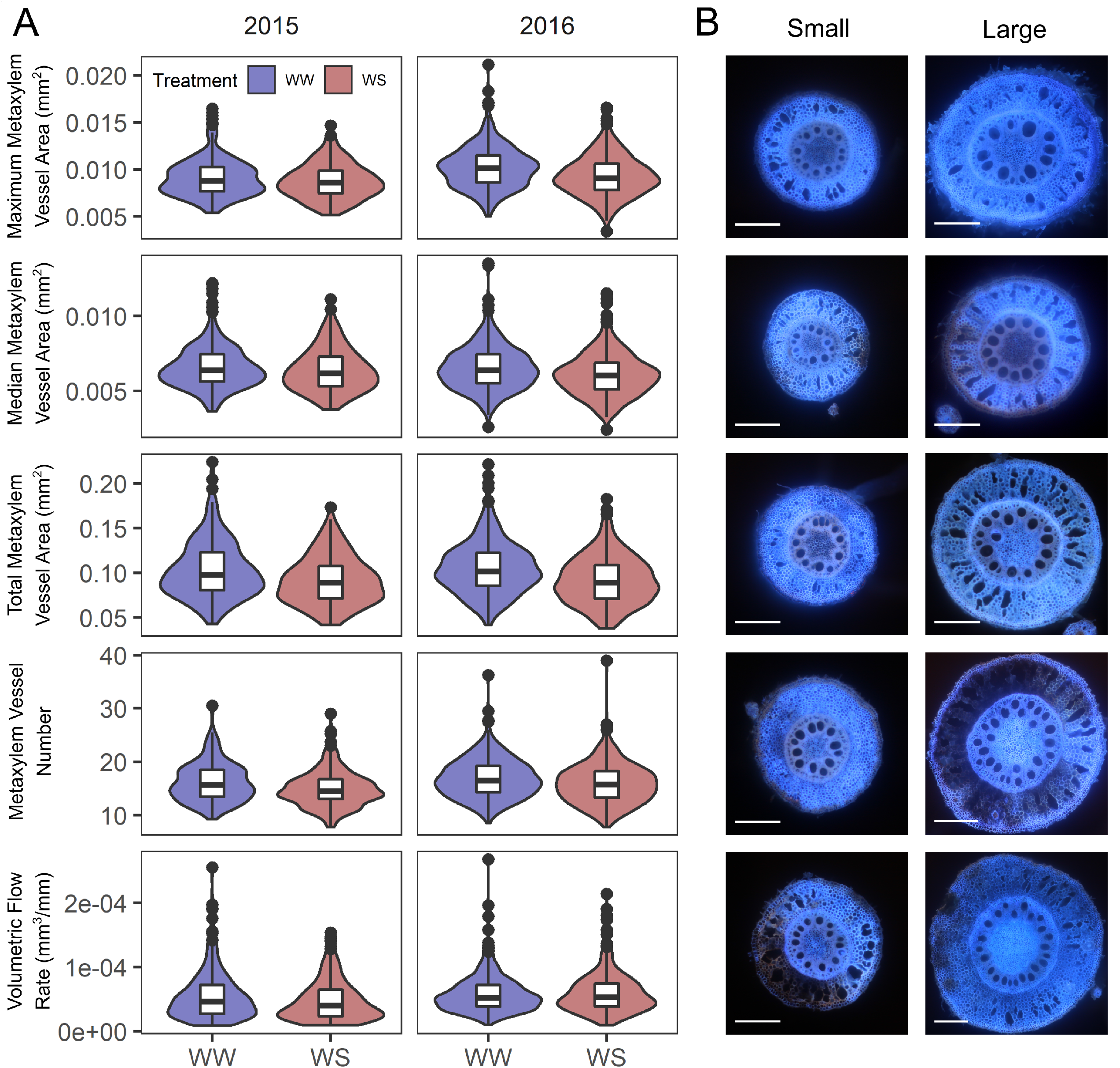
Wide variation exists in metaxylem phenes under well-watered and water stress conditions in the field. (A) Violin plots show the distribution of each metaxylem phene under well-watered (WW, blue) and water stress (WS, red) in each field season while the overlaid box plots show the median bounded by the first and third quartile. Two-way ANOVA determined statistical differences between treatments and growth seasons (Table 1). (B) Root cross-sectional images collected via laser ablation tomography illustrate the range in each metaxylem phenotype. White scale bars in the bottom left of each image are equal to 0.5 mm.

**Table 1.**
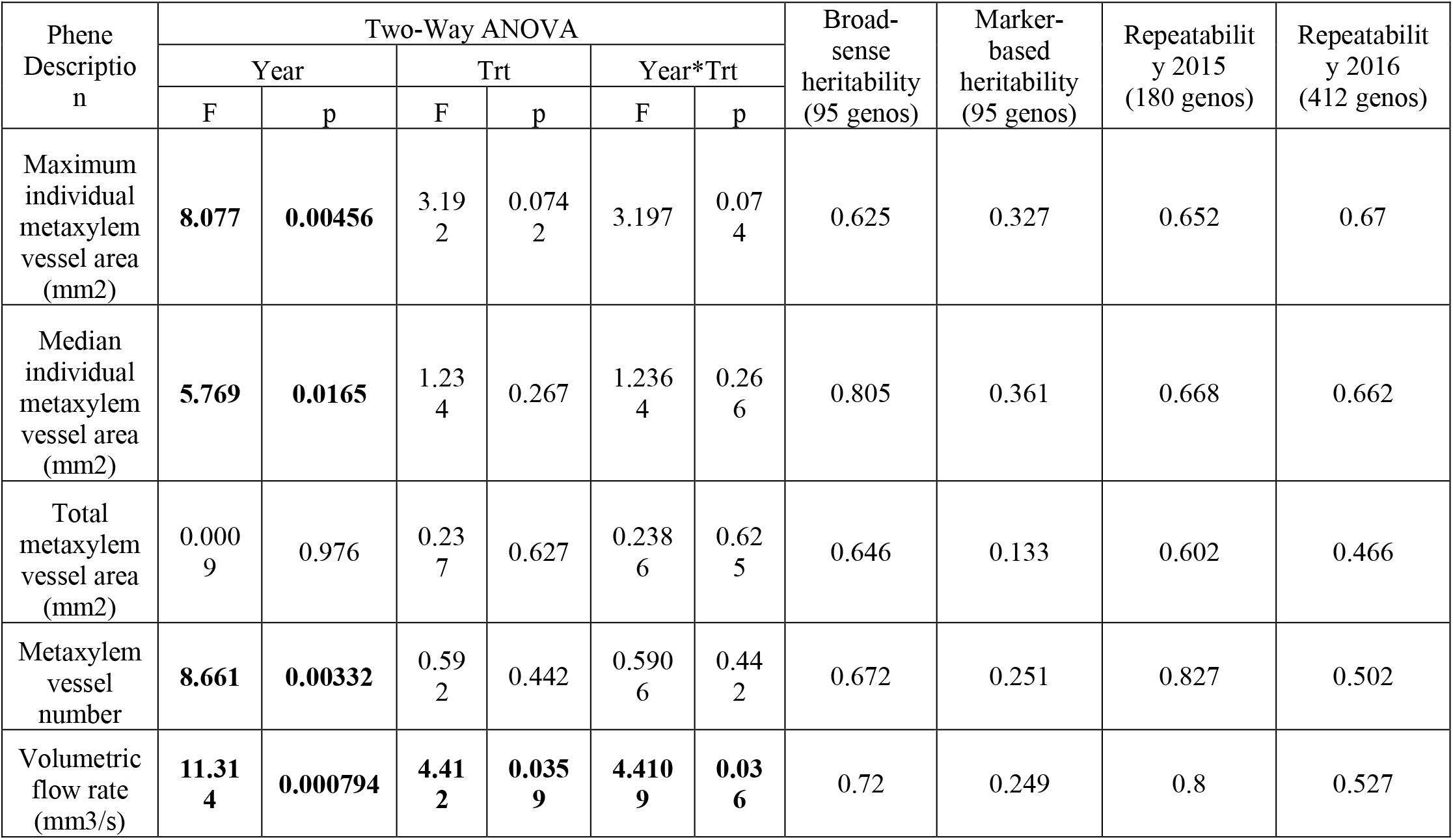
Two-way ANOVA of root metaxylem phenes measured across two field seasons under well-watered (WW) and water stress (WS) conditions, and broad-sense and marker-based heritabilities calculated for each phene among 95 genotypes planted in each season. Repeatability of each phene was calculated using all genotypes grown in each season. Bold numbers in the two-way ANOVA indicate significant associations (*p* < 0.05).

| Phene<br>Descriptio<br>n | Two-Way ANOVA |  |  |  |  |  | Broad-sense<br>heritability<br>(95 genos) | Marker-based<br>heritability<br>(95 genos) | Repeatabilit<br>y 2015<br>(180 genos) | Repeatabilit<br>y 2016<br>(412 genos) |
| --- | --- | --- | --- | --- | --- | --- | --- | --- | --- | --- |
|  | Year |  | Trt |  | Year*Trt |  |  |  |  |  |
|  | F | p | F | p | F | p |  |  |  |  |
| Maximum<br>individual<br>metaxylem<br>vessel area<br>(mm2) | **8.077** | **0.00456** | 3.19<br>2 | 0.074<br>2 | 3.197 | 0.07<br>4 | 0.625 | 0.327 | 0.652 | 0.67 |
| Median<br>individual<br>metaxylem<br>vessel area<br>(mm2) | **5.769** | **0.0165** | 1.23<br>4 | 0.267 | 1.236<br>4 | 0.26<br>6 | 0.805 | 0.361 | 0.668 | 0.662 |
| Total<br>metaxylem<br>vessel area<br>(mm2) | 0.000<br>9 | 0.976 | 0.23<br>7 | 0.627 | 0.238<br>6 | 0.62<br>5 | 0.646 | 0.133 | 0.602 | 0.466 |
| Metaxylem<br>vessel<br>number | **8.661** | **0.00332** | 0.59<br>2 | 0.442 | 0.590<br>6 | 0.44<br>2 | 0.672 | 0.251 | 0.827 | 0.502 |
| Volumetric<br>flow rate<br>(mm3/s) | **11.31**<br>**4** | **0.000794** | **4.41**<br>**2** | **0.035**<br>**9** | **4.410**<br>**9** | **0.03**<br>**6** | 0.72 | 0.249 | 0.8 | 0.527 |

Broad-sense and marker-based heritability on an entry mean basis was calculated within a shared set of 95 genotypes. Broad-sense heritability ranged from 0.625 to 0.805 and marker-based heritability ranged from 0.133-0.361, overall (Table 1). Repeatability was also calculated for the collection of genotypes grown in each season and ranged from 0.602-0.827 in 2015 and 0.466-0.670 in 2016 (Table 1).

### GWAS for metaxylem phenes

A panel of 899,794 SNPs (Mazaheri et al., 2019) was used for GWAS on all root metaxylem phenes under well-watered and water-stress conditions conducted with GAPIT (Lipka et al., 2012). The fit of the model was corroborated with quantile-quantile (Q-Q) plots (Supplemental Figure S2). Separate GWAS analyses were performed for each treatment and season of field phenotype data to identify genetic loci that consistently exhibited similar effects on variance in root metaxylem phenotypes. Despite differences in taxa membership in each season, there were 550,617 SNPs included in analyses of each season that were used to evaluate the consistency of minor allele effects on phenotypes (Supplemental Figure S3A). Few of the SNPs omitted from the comparison were significant in any analysis. Because of its larger genotype collection, the 2016 GWAS was designated the “testing set” to identify many genetic loci potentially associated with root metaxylem phenotypes. The 2015 GWAS was the “validation set” to corroborate allelic effects observed in the testing set. For the testing and validation sets, the allelic effect of each SNP in each phene was converted to a percentile rank where 0 denoted a strong negative effect and 100 indicated a strong positive effect on phenotype. A SNP with consistent allelic effects for the same phene was one where the percentile rank in the validation set was within 10 percentile points of its rank in the testing set (Supplemental Figure S3). This analysis was repeated for each root metaxylem phene and treatment. The number of SNPs that displayed consistent allelic effects was 24-28% of the full SNP panel included in each GWAS and varied among phenes and treatments (Supplemental Table S2). A SNP was significant if it exceeded the p-threshold and displayed consistent allelic effects on metaxylem phenotypes. No significant associations were detected above the Bonferroni threshold of -log(*p*) = 9.08. Thus, a more liberal threshold of -log(*p*) = 3.43 was applied to accommodate SNPs associated with small effect sizes (Figure 2). This threshold was estimated using a model that took into account the average marker-based heritability of all root metaxylem phenes and linkage disequilibrium in maize (Kaley and Purcell 2019).

**Figure 2.**
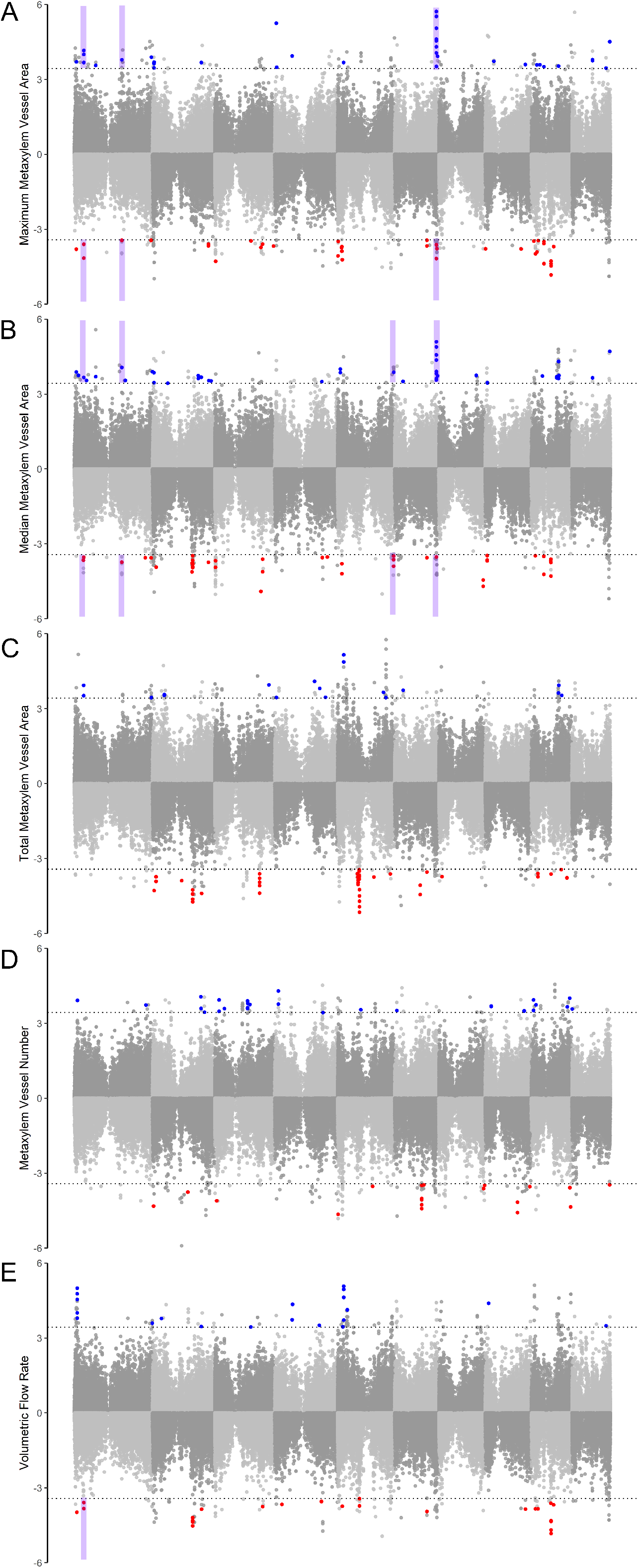
Manhattan plots of the results of the GWAS for (A) maximum metaxylem vessel area, (B) median metaxylem vessel area, (C) total metaxylem vessel area, (D) metaxylem vessel number and (E) volumetric flow rate in each watering regime in 2016 (grey). Results from the well-watered studies are shown on the positive axis while the water stress is on the negative axis. The dotted line indicates the significance threshold (-log(*p*) > 3.43). Dots colored blue and red are significant SNPs with consistent allelic effects in the 2015 GWAS in the well-watered and water-stressed treatments, respectively. Select genomic regions that shared significant SNPs in both treatments have been highlighted in purple.

Overall, 244 gene models contained or were within 5kb of 351 significant SNPs associated with maize root metaxylem phenes under well-watered and water stress conditions (Table 2, Figure 2, Supplemental Table S3). Under well-watered conditions, 181 significant SNPs located in or near 124 gene models were associated with metaxylem phenes (Table 2, Supplemental Table S3). Under water stress, 176 significant SNPs located in or near 126 gene models were associated with metaxylem phenes (Table 2, Supplemental Table S3). Several genomic regions displayed a high density of significant SNPs associated with a specific phene (Figure 2). For instance, 13 gene models near 25 significant SNPs associated with total metaxylem vessel area under water stress were detected in a region on chromosome 5. Another region on chromosome 3 contained 5 significant SNPs nearby two candidate gene models also associated with total metaxylem vessel area under WS. Two gene models nearby 14 significant SNPs associated with median metaxylem vessel area under water stress were detected on chromosome 2. Seven gene models near 12 significant SNPs associated with maximum and/or median metaxylem vessel area under well-watered conditions were detected in a region on chromosome 6. Of the candidate genes associated with root metaxylem phenes under well-watered and water stress, approximately 68% were annotated for Mapman ontogenic categories (Supplemental Figure S4). However, there were few differences in representation of Mapman ontogenic categories for candidate genes associated with the well-watered treatment compared to those associated with water stress. A slightly larger proportion of candidate genes identified under water stress were associated with developmental processes while a larger proportion of candidate genes identified under well-watered conditions were associated with stress responses.

**Table 2.**
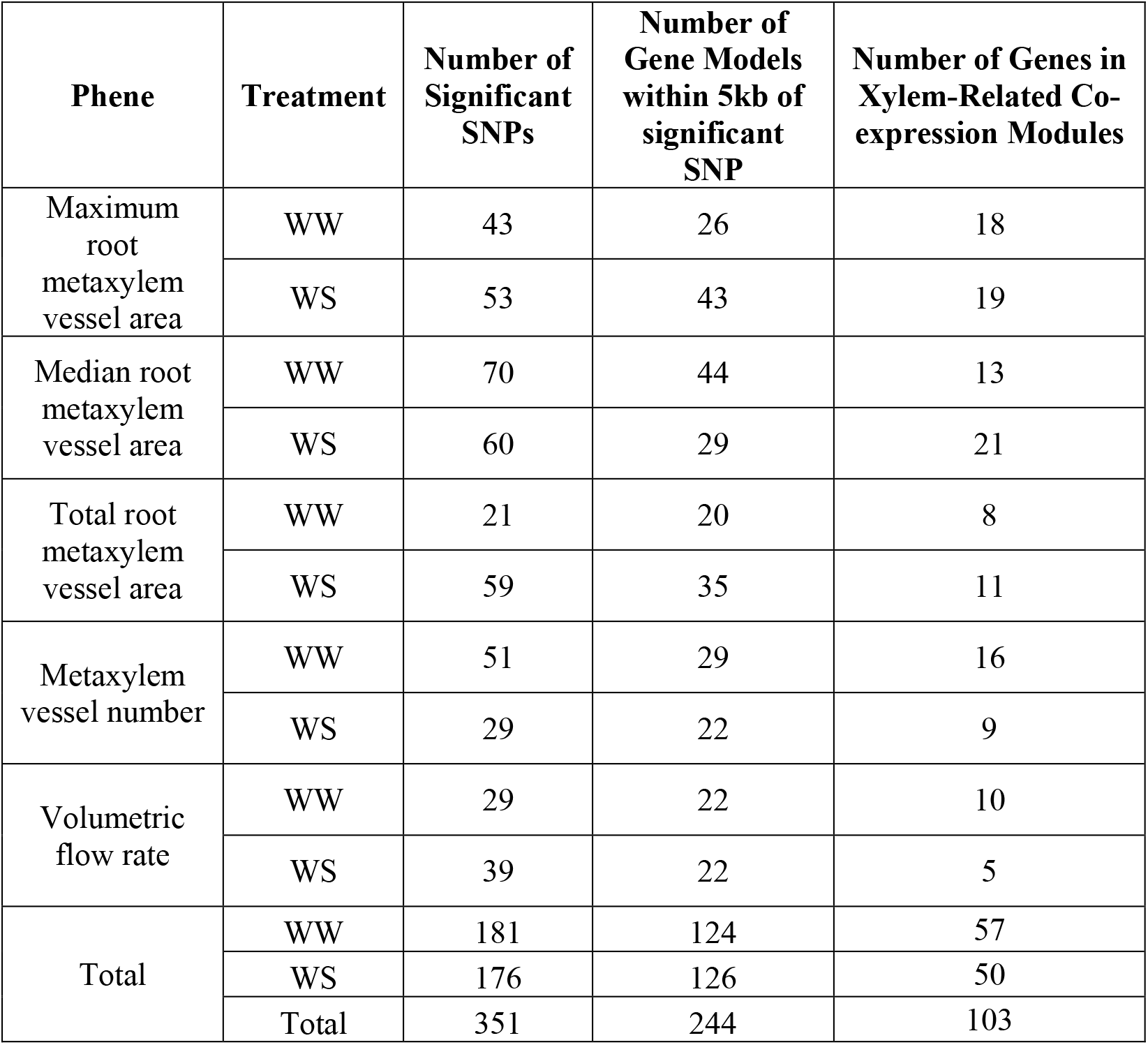
Number of significant SNPs and candidate genes detected in the maize GWAS and the number of candidate genes found in one of 14 gene co-expression modules associated with root tissues most relevant to xylem development.

The Variant Effect Predictor tool was used to assess the likelihood of the significant SNPs being the causal SNP for mutation in the gene model (McLaren et al., 2016). In well-watered conditions and water stress, approximately 35% and 22% of variants, respectively, generated missense or 5’ UTR variants, which are likely to affect protein function or transcriptional regulation (Figure 3). Significant SNPs resulting in missense mutations were over three times as prevalent in candidate genes associated with well-watered conditions (26.7% of all variants) compared to water stress (7.7% of all variants). In both watering regimes, approximately 17% of the SNPs resulted in synonymous mutations likely having no effect on protein function. Nonsense mutations resulting in a premature stop codon were extremely rare.

**Figure 3.**
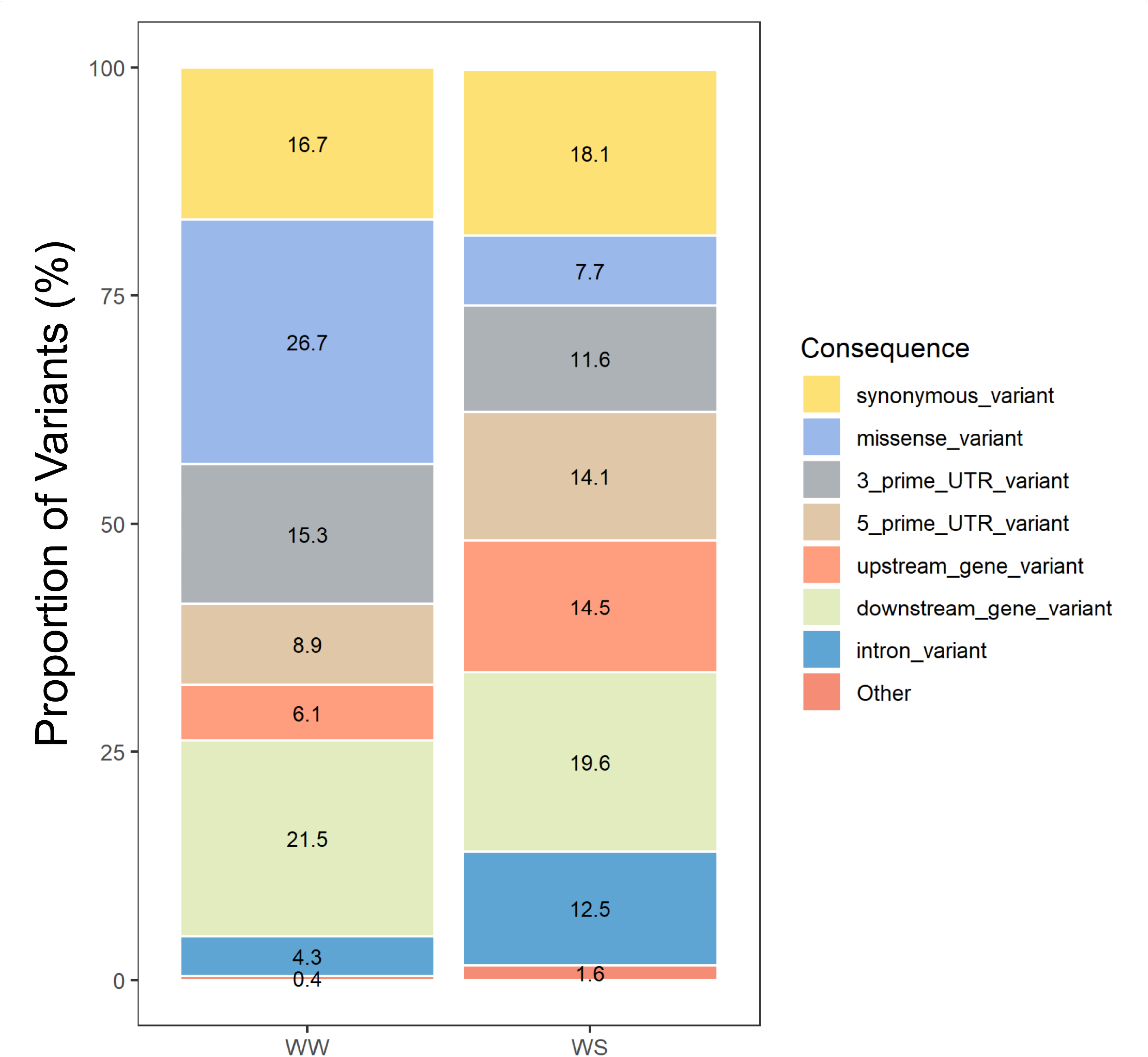
Predicted consequences of minor allele variants presented as a proportion of the total number of significant SNPs in well-watered (WW) and water stressed (WS) conditions.

We found 13 significant SNPs associated with six candidate genes co-localized in both watering regimes (Figure 2, Supplemental Table S4). Several of these candidate genes also co-localized in more than one metaxylem phene either by association with a single co-localized SNP or because multiple SNPs residing near the gene were detected. Of these candidate genes, only one was annotated. This gene was Zm00001d018394, DNA-binding protein phosphatase 2C (DBPTF2), and contained four SNPs significantly associated with median root metaxylem vessel area in both water treatments. The other five genes are not annotated, but their functional annotation may be inferred according to its homology with genes characterized in *Arabidopsis* (Figure 2, Supplemental Table S4). The gene Zm00001d028486 encodes a chemocyanin and contains two SNPs associated with median and maximum root metaxylem vessel areas in both treatments and volumetric flow rate under water stress. Zm00001d031332 encodes a heat shock protein and contains four SNPs associated with median and maximum root metaxylem vessel areas under well-watered and water stress conditions. Two SNPs nearby Zm00001d038573, a ubiquitin domain containing 1, and Zm00001d038754, an endoplasmic reticulum lumen protein retaining receptor, were associated with maximum and median root metaxylem vessel area in both treatments. Finally, the gene Zm00001d045954 encodes a RNA binding protein that contains one SNP significantly associated with maximum root metaxylem vessel area in both treatments.

### Gene Co-expression Network of root tissues

To identify groups of genes with a high likelihood of regulating root xylem phenotypes, the WGCNA package (Langfelder and Horvath, 2008) was used to discover gene co-expression relationship networks among expressed genes in 19 root tissues (Stelpflug et al., 2016) (Figure 4). Using a signed network analysis, 39 modules of co-expressed genes were identified, each one labeled as a color (Figure 4A). The membership of each module ranged from 35 (“lavenderblush3”) to 5070 (“black”) genes. Of those, 14 modules had greater potential for being associated with metaxylem phenotypes because of the significant correlation between the module eigengene and root tissue samples relevant to xylem development (Figure 4B). Root tissues most relevant to xylem development (-log(*p*) > 2) are the stele 3 days after sowing (DAS) (modules “lightcyan”, “thistle2”, “yellowgreen”), the differentiation zone 3 DAS (“darkred”, “lightcyan1”, “red”), the meristematic and elongation zone 3 DAS (“black”, “lightgreen”, “thistle1”), the whole root system 3 DAS (“salmon4”), zone 1 of the primary root 7 DAS (“mediumpurple3”, “orange”), and zone 2 of the primary root 7 DAS (“darkgreen”, “navajowhite2”). Gene ontology (GO) analysis showed that several biological functions related to xylem development were over-represented in the xylem-related modules (Figure 4C). Modules “black” and “lightgreen” were most strongly associated with root morphology and anatomical development whereas module “thistle2” was particularly associated with the cell wall and carbohydrate biosynthesis. GO terms related to signaling were most strongly represented in modules “lightcyan”, “lightgreen” and “yellowgreen”. While the “black” module was most strongly associated with stress responses, six additional modules were shown to be associated with stress responses as well. No GO terms were over-represented by modules “lightcyan1”, “thistle1” and “salmon4” (not shown).

**Figure 4.**
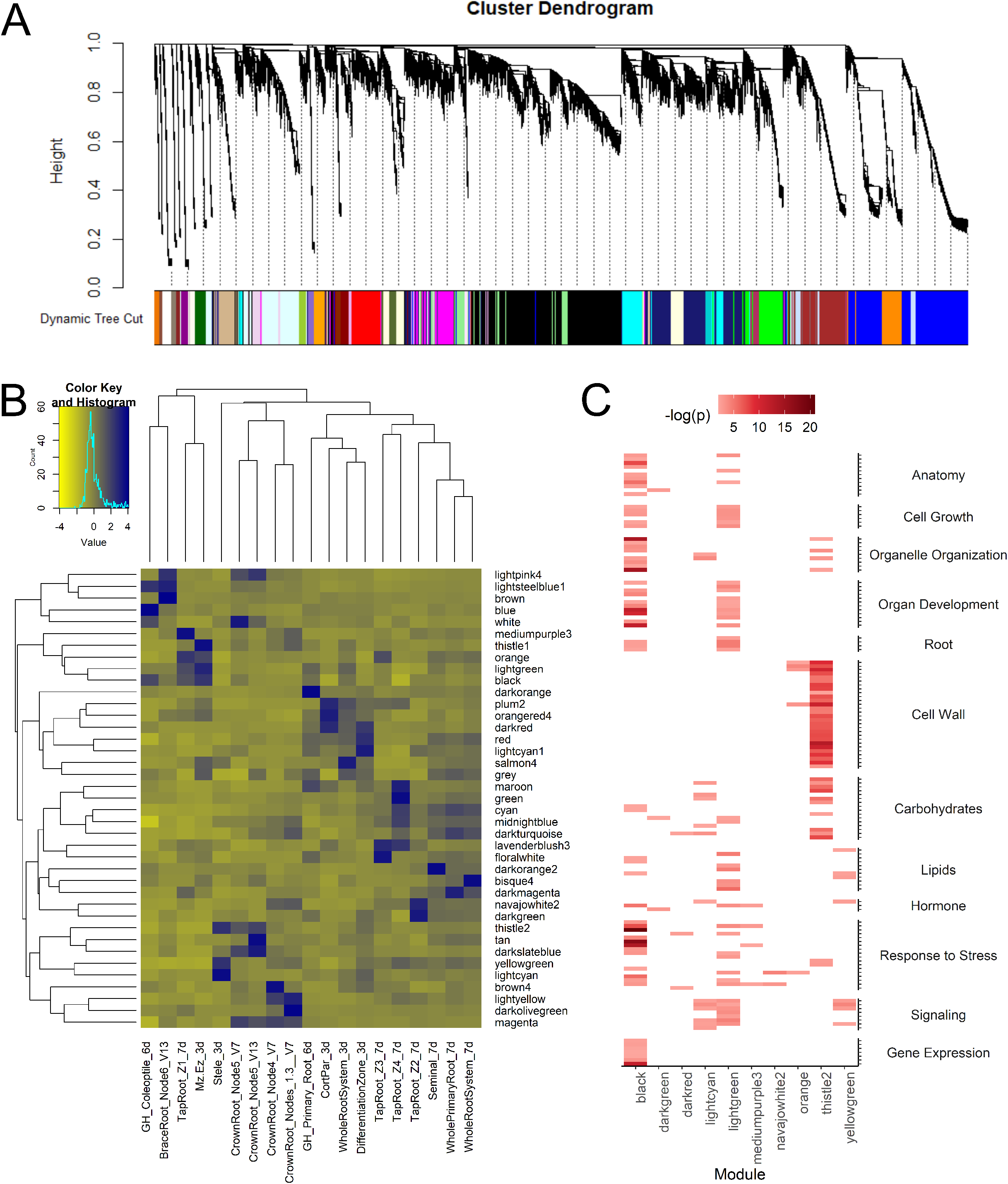
A gene co-expression network and significant correlation of modules. (A) Hierarchical clustering dendrogram displaying 39 modules of co-expressed genes. (B) A heatmap showing the significance of the correlation (-log(*p*)) between modules and various root tissues. (C) Inferred biological function of modules most likely associated metaxylem development based on GO analysis. Only GO terms most relevant to xylem development are displayed. No GO terms were over-represented by modules “lightcyan1”, “salmon4”, and “thistle1”.

Of the 244 significant gene candidates identified in the GWAS, 103 reside in gene co-expression modules relevant to xylem development – 57 under well-watered conditions and 50 under water stress (Table 2). Topology metrics of each module’s network were used to infer the essentiality of each gene within the network (Supplemental Table S5). Hub genes (HUB), or those that were most connected to the rest of the network, exhibit a high closeness centrality while bottleneck genes (BN), those through which many connections are channeled, exhibit a high betweenness centrality. A gene may be both a bottleneck and hub (BN/HUB). Within the network of each co-expression module, genes in the 80th percentile and above of closeness centrality and betweenness centrality were classified as hub and bottleneck genes, respectively (Supplemental Figures S5). Of the GWAS candidate genes that reside in stele-related modules (“lightcyan”, “thistle2”, “yellowgreen”), 3 hub genes, 1 bottleneck genes, and 3 bottleneck/hub genes were identified (Figure 5). Only one of these genes is annotated: ferritin homolog2 (FER2; Zm00001d023225), a hypothetical bottleneck/hub gene from “thistle2” associated with root metaxylem vessel number under well-watered conditions (Figure 5)..

**Figure 5.**
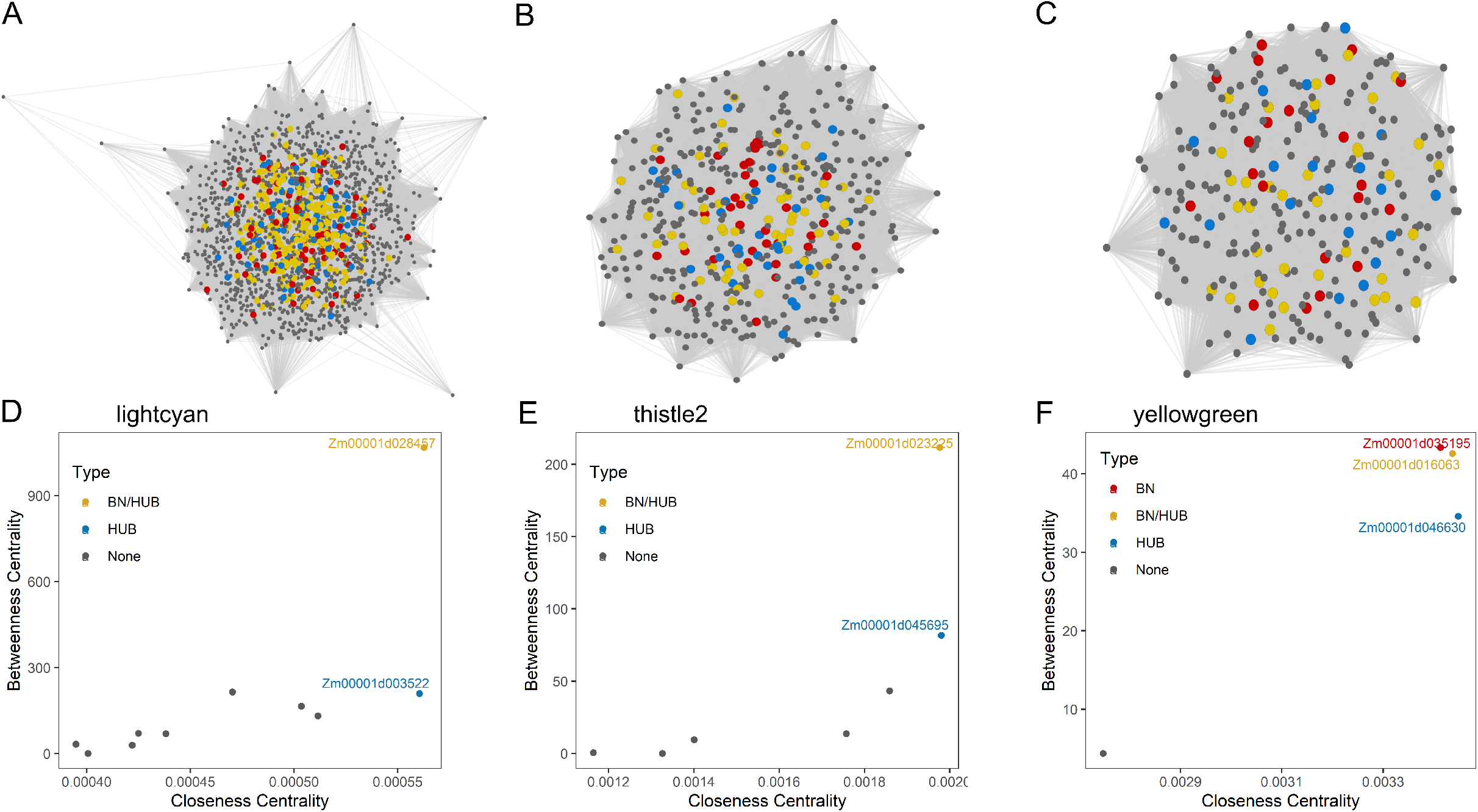
Gene co-expression subnetworks for modules associated with root stele 3 days after sowing. These modules were “lightcyan” (A, D), “thistle2” (B, E) and “yellowgreen” (C, F). Network visualizations (A, B, C) show all interactions within the subnetwork. Scatterplots (D, E, F) show the calculated closeness centrality and betweenness centrality for candidate genes determined by GWAS residing in the corresponding module. All bottleneck (BN), bottleneck/hub (BN/HUB) and hub (HUB) genes are shown in red, yellow, and blue, respectively.

### Comparative GWAS on metaxylem phenes between maize and rice

Because the development of metaxylem phenotypes in maize and rice is similar, we hypothesized that these species have shared genetic control for metaxylem phenotypes. To test this hypothesis, we compared the unique candidate genes from each maize and rice GWAS to identify syntenic genes that were similarly significant in each analysis. We identified eight pairs of syntenic genes in maize and rice that were significantly associated with metaxylem phenes in a GWAS: six were related to maize grown under well-watered conditions and two were related to maize grown under water stress (Figure 6A, Table 3). Five of these syntenic pairs were similarly associated with size-related metaxylem phenes (maximum, median and total metaxylem vessel areas; volumetric flow rate) or vessel number in both maize and rice. Some of these genes may share similar functional roles since several metabolic pathways were represented by at least two syntenic pairs. Two maize genes (Zm00001d008569, Zm00001d046838) and their respective rice homologs (Os01g0134500, Os06g0334300) are active in the brassinosteroid biosynthesis pathway and were each associated with metaxylem size-related phenes under well-watered conditions (Table 3). Four of these syntenic gene pairs are associated with drought responses. Zm00001d038999 encodes the maize protein drought-induced 19 and was associated with maximum and median vessel area in maize under well-watered conditions, while its rice homolog (Os05g0562200) was associated with metaxylem vessel number in the *indica* rice subpopulation. Zm00001d048165 and its rice homolog Os03g0353400 encode the protein EARLY RESPONSIVE TO DEHYDRATION 15, which was similarly associated with metaxylem vessel number in the full rice population and maize under WS. However, there were no significant associations between the minor allele and root metaxylem phenotypes observed in maize or rice for either of these genes.

**Figure 6.**
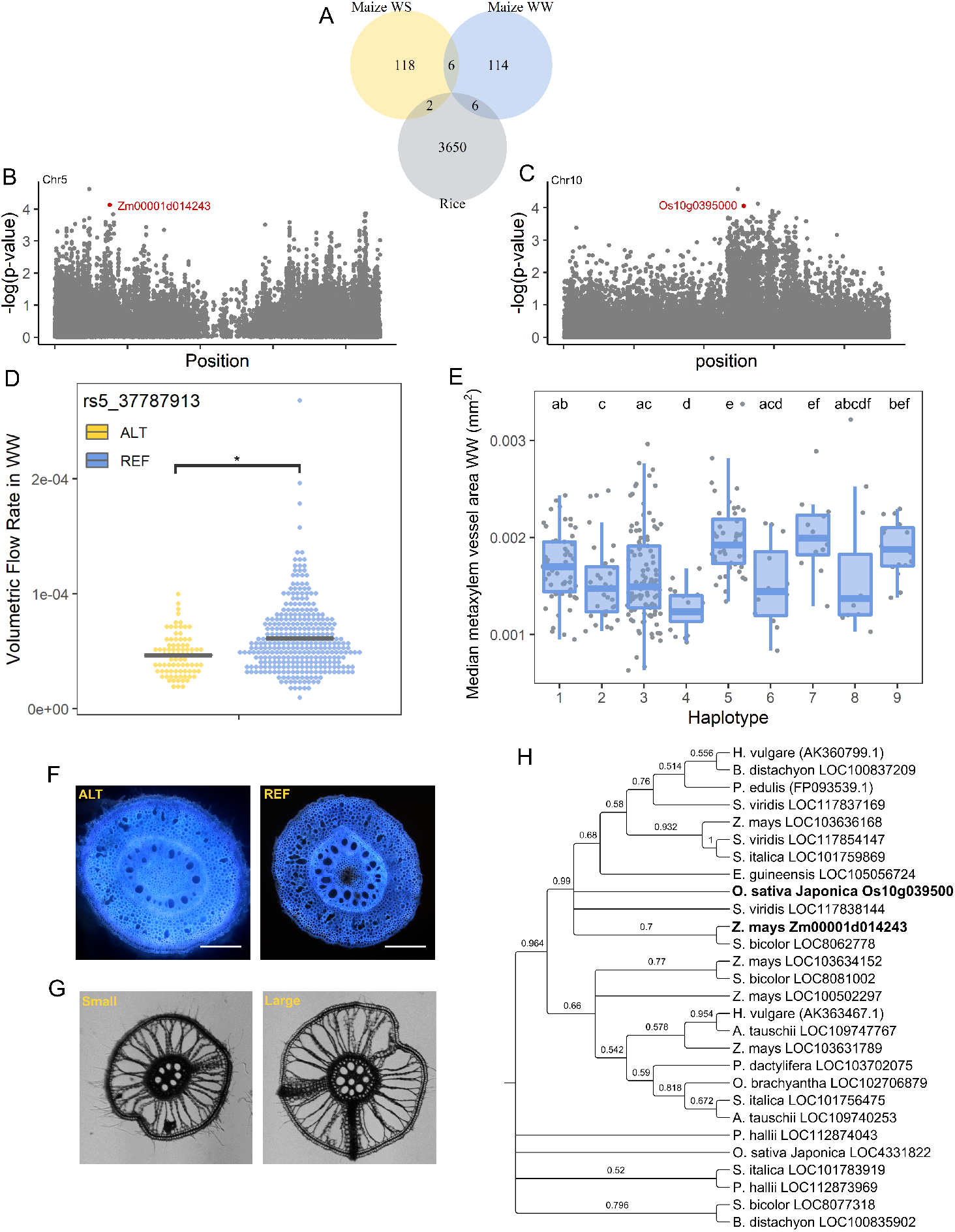
Comparative GWAS between maize and rice for metaxylem phenes identified a syntenic gene pair encoding root-specific kinase 1 associated with root metaxylem phenotypes. (A) Comparison of the results of three GWAS seeking genetic loci associated with root metaxylem phenes in maize under well-watered (WW) and water-stressed (WS) conditions and rice. (B) Manhattan plot of SNPs and their associated with the volumetric flow rate under WW on maize chromosome 5 highlighting the candidate gene (Zm00001d014243) in red. (C) Manhattan plot of SNPs associated with metaxylem vessel area on rice chromosome 10 highlighting the candidate gene (Os10g0395000) in red. (D) Significant differences in volumetric flow rate in WW are associated with the minor allele (ALT, yellow) compared to the major allele (REF, blue) determined by two-sample *t*-test (*p* < 0.001). (E) Significant differences in metaxylem vessel area are associated with nine haplotypes of Os10g0395000 determined by Kruskal-Wallis (*p* < 0.001). Haplotypes are disclosed in Supplemental Figure S6. (F) Representative images maize root cross-sections captured via LAT illustrating visual phenotypic differences in individuals that contained the minor (ALT) and major (REF) alleles. (G) Representative images of rice root cross-sections collected by hand-sectioning illustrating visual contrasts in metaxylem vessel area. (H) Phylogenetic tree of gene homologous to root-specific kinase 1. The amino acid sequences of 28 proteins of high sequence similarity were aligned by MUSCLE and the phylogenetic tree was constructed using MEGA version 10.1.8 and TreeGraph 2. Bootstrap values from 500 replicates were used to assess the robustness of the tree. The maize and rice candidate genes identified in GWAS are labeled in bold.

**Table 3.** Syntenic gene pairs that were strongly associated with root metaxylem phenes in both maize and rice.

| Maize Gene | Maize Root Phene | Rice Gene | Rice Root Phene | Rice Subpopulation | Description |
| --- | --- | --- | --- | --- | --- |
| Zm00001d008569 | MXAtot_WW, VFR_WW | Os01g0134500 | MXAmed | IND | Delta(7)-sterol-C5(6)-desaturase 1 |
| Zm00001d011147 | MXAmax_WS | Os09g0570400 | MXN | ALL | Ascorbate transporter, chloroplastic |
| Zm00001d014243 | VFR_WW | Os10g0395000 | MXAmed | AUS | root-specific kinase 1 |
| Zm00001d017216 | MXAtot_WW | Os02g0617700 | MXAmed | IND | Polyribonucleotide nucleotidyltransferase 2, mitochondrial |
| Zm00001d038999 | MXAmax_WW, MXAmed_WW | Os05g0562200 | MXN | IND | drought-induced 19 |
| Zm00001d039000 | MXAmax_WW, MXAmed_WW | Os05g0562300 | MXN | IND | HSP40/DnaJ peptide-binding protein |
| Zm00001d046838 | MXAmed_WW, MXAtot_WW | Os06g0334300 | MXAmed | ALL | Putative receptor-like protein kinase family protein |
| Zm00001d048165 | MXN_WS | Os03g0353400 | MXN | ALL | EARLY RESPONSIVE TO DEHYDRATION 15 (ERD15) |

A root-specific kinase 1, Zm00001d014243 and its rice homolog Os10g0395000, were similarly associated with metaxylem size-related phenes under well-watered conditions: volumetric flow rate in maize and median root metaxylem vessel area in the *aus* rice subpopulation (Figure 6). In maize, individuals containing the minor allele exhibited on average a significantly lower volumetric flow rate compared to the rest of the population (p < 0.001; Figure 6). Substitution with this minor allele results in a missense mutation leading to a replacement of serine for threonine. Because it was distant from the associated significant SNP in rice, phenotypic differences were evaluated for various haplotypes of the gene candidate (Supplemental Figure S6). There were significant differences (*p* < 0.001) in root metaxylem vessel area among haplotypes where individuals with haplotype 4 contained the smallest root metaxylem vessel area and those of haplotype 7 contained the largest (Figure 6). BLAST was used to collect highly similar sequences of this gene in other species. This syntenic gene pair exhibited a high degree of homology with root specific kinases and serine/threonine protein kinases of several other monocotyledonous species (Figure 6). Root-specific kinase 1 also has several homologs in maize and rice. No sequences of dicots were identified under these parameters.

## Discussion

Since metaxylem anatomy has important implications for hydraulic function (Hacke and Sperry, 2001; Comas et al., 2013), the genetic architecture of root metaxylem phenotypes has important implications for plant performance under drought. We identified several candidate loci associated with root metaxylem phenes under well-watered and water stressed conditions by integrating a GWAS conducted in maize with a root-specific gene co-expression network and by performing a cross-species comparative GWAS with rice. We observed wide variation in root metaxylem phenotypes in maize, and this variation is under distinct genetic control in optimal and water-limited environments. We observed high heritability in root metaxylem phenes (Table 1). Broad-sense heritability observed was higher that previously reported values for root anatomical phenes (Kadam et al., 2017; Schneider et al., 2020a), but may be slightly conflated given that these values were based on only 95 genotypes than were planted in each season.Mean values of all metaxylem phenes apart from volumetric flow rate were unchanged in response to drought stress (Figure 1), but responses to drought varied by genotype (Supplemental Figure S1). There were few genetic loci associated with root metaxylem phenes that co-localized between water regimes (Supplemental Table S4, Figure 2, Figure 6A), which suggests that drought induces changes in vascular patterning distinctly different from the processes employed when water is more readily available. Unexpectedly, three times as many non-synonymous consequences created by minor allele substitutions were associated with root metaxylem phenes in the well-watered treatment compared to water stress (Figure 3). Modifications in plant function due to a missense minor allele substitution may be more impactful under water stress because of the greater selection pressure acting on these alleles compared to well-watered conditions. The mechanisms counteracting potential functional impairments due to missense mutation are perhaps more robust under water stress (Balestrazzi et al., 2011), leading to reduced prevalence of missense variants compared to non-stressed conditions.

Our GWAS results identified several candidate genes associated with root metaxylem phenes under well-watered and water stress conditions, and these candidate genes likely serve functional roles relevant to xylem development according to the gene co-expression network. GWAS has been widely utilized as a tool to identify genes related to a particular phenotype, but it also presents many challenges. First, GWAS identifies many genes of which a notable proportion are false positives. Instead of relying solely on significance values associated with SNPs, we sought candidate genes with strong and consistent associations with root metaxylem phenotypes in maize across two field seasons and in a cross-species GWAS comparison with rice. Second, phenotyping highly complex quantitative traits associated with small effect sizes remains a challenge in GWAS (Ingvarsson and Street, 2011; Schneider et al., 2020a; Schneider et al., 2020b), which in turn makes candidate gene selection difficult. Integrating GWAS with a gene co-expression network enables candidate gene selection to be determined by network relationships inferring gene essentiality and biological function in its respective module (Yu et al., 2004; Yu et al., 2007; McDermott et al., 2009). Of the 103 candidate genes identified through GWAS that belonged to gene co-expression modules associated with root tissues most relevant to xylem development, we identified 7 bottleneck genes, 9 hub genes, and 17 bottleneck/hub genes (Figure 5, Supplemental Figure S5). We argue that these candidate genes that serve as hub and/or bottleneck genes within a gene co-expression network merit greater attention for validation in functional studies. Because these genes are more essential for maintaining gene-gene interactions and signaling cascades, mutation of hub and bottleneck genes may have a greater chance of inducing lethality. However, they also promise a higher likelihood of producing substantial modification in phenotype and function.

Because of their contribution to xylem vessel wall development and reinforcement, genes involved in cell wall processes are natural targets for further investigation. Three genes in xylem-related co-expression modules are associated with cell wall-related processes and lignin biosynthesis according to their Mapman annotation: alpha-L-arabinofuranosidase /beta-D-xylosidase isoenzyme ARA-I (Zm00001d009060) associated with cell wall degradation, pectinesterase (Zm00001d022567), and 4-coumarate--CoA ligase-like 4 (Zm00001d027519) (Supplemental Table S3). Four additional candidate genes were proposed to be associated with the biosynthesis and metabolism of cell-wall polysaccharides (Okekeogbu et al., 2019) (Supplemental Table S3): a pectate lyase (Zm00001d015328), a chitin elicitor-binding protein (Zm00001d027533), a protein disulfide isomerase (Zm00001d046399), and a glucan endo-1,3-β-glucosidase (Zm00001d047712). Interestingly, all candidate genes in this group were associated with root metaxylem size-related phenes under water stress. Surprisingly, we did not find any significant associations with cell wall-related genes that are well known to directly influence xylem development. For instance, cellulose synthase 4 (CesA4), which contained a SNP with a consistent effect on phenotype that was just below the p-threshold (-log(*p*) = 3.02), is a known positive regulator of xylem differentiation in *Arabidopsis* (Taylor et al., 2003) and in turn is regulated by VASCULAR-RELATED NAC DOMAIN7 (VND7) (Yamaguchi et al., 2011). Much research has demonstrated that VND6 and VND7 are key regulators of xylem vessel differentiation in *Arabidopsis* (Kubo et al., 2005; Yamaguchi et al., 2010a; Yamaguchi et al., 2010b; Yamaguchi et al., 2011; Turco et al., 2019), but neither of these genes have yet been characterized in maize. Maize homologs of VND6 – NAC10, NAC27, NAC46, NAC131 – and VND7 – NAC62 and NAC92 – were co-expressed in the three modules strongly associated with stele tissues 3 DAS (“lightcyan”, “thistle2”, “yellowgreen”), but none of these genes were associated with significant SNPs detected in GWAS.

Several gene candidates associated with phytohormone biosynthesis and signaling were identified in our GWAS analyses. Brassinosteroid-deficient *Arabidopsis* mutants were severely dwarfed, exhibited impaired root elongation, and produced fewer xylem in leaf vascular bundles (Szekeres et al., 1996; Choe et al., 1999a; Choe et al., 1999b). In our cross-species GWAS, two additional gene candidates involved in brassinosteroid metabolism were associated with various metaxylem vessel area-related phenes under WW in both maize and rice (Table 3). Δ(7)-sterol-C(5)-desaturase (Zm00001d008569, Os01g0134500) is the rate-limiting step in brassinolide biosynthesis and impairment of its activity is linked to defective longitudinal growth, irregularly spaced vascular bundles, and reduced xylem vessel size and number (Choe et al., 1999b). An additional gene syntenic pair (Zm00001d046838, Os06g0334300) encoding a putative receptor-like kinase family protein sharing a high degree of homology to HERCULES1 (HERK1), which is regulated by brassinosteroids and required for cell elongation (Guo et al., 2009a; Guo et al., 2009b). The cross-species GWAS comparison also identified the syntenic gene pair for a DnaJ/heat shock protein 40 family protein (Zm00001d039000/Os05g0562300) that was significantly associated with maximum and median metaxylem vessel areas under well-watered conditions and metaxylem vessel number in the rice *indica* subpopulation. Interestingly, a member of this family has been shown to mediate environmental stress responses via brassinosteroid signaling – over-expression of the gene maintained proper cell elongation in the inflorescence and roots in the presence of a brassinosteroid inhibitor (Bekh-Ochir et al., 2013). In the maize GWAS, five additional genes with potential associations with hormone metabolism and biosynthesis were identified (Supplemental Table S3). A flavonol synthase (Zm00001d002100) and AP2/EREB transcription factor 209 (EREB209) are involved in ethylene metabolism and were significantly associated with total root metaxylem vessel area and volumetric flow rate, respectively, under water stress. Ethylene has been shown in *Zinnia elegans* cultures to play a dual role by stimulating the rate of tracheary element differentiation and controlling the stem cell pool size during secondary xylem formation (Pesquet and Tuominen, 2011). Auxin response factor 29 (ARF29) and aux/IAA-transcription factor 22 (IAA22) were significantly associated with various root metaxylem vessel size related phenotypes under water stress and well-watered conditions, respectively. Auxin is a key regulator of nearly every plant developmental process including many aspects of vascular development (Milhinhos and Miguel, 2013). Most recently, auxin application was found to repress transcription of NACs necessary for fiber and secondary cell wall development and promote vessel-specific NACs in *Populus* stems (*Johnsson et al., 2019).* The function of these gene candidates has been mostly characterized in shoots, but our results suggest they may also contribute to variation in root metaxylem phenotypes.

Interestingly, the gene encoding a nodulin-like family protein (Zm00001d005082) contained 11 significant SNPs associated with maximum, median, and total metaxylem vessel areas and volumetric flow rate under water stress and co-expressed with genes highly expressed in the root meristematic/elongation zone 3 DAS (Supplemental Table S3). This nodulin lies within the genomic region on chromosome 2 that was densely populated with SNPs strongly associated with total root metaxylem vessel area under water stress and may also be a hub gene since its closeness centrality was relatively high (Supplemental Figure S5). Another nodulin protein (Zm00001d037160) contained one SNP strongly associated with metaxylem vessel number and co-expressed with genes highly expressed in the stele 3 DAS (Supplemental Table S3). Nodulins are not specific to legumes since homologs of nodulins have been found in several non-nodulating species (Denancé et al., 2014). In non-nodulating species, nodulin-like proteins are transmembrane proteins shown to transport a wide range of compounds and one has been shown to enhance root hair elongation in rice (Huang et al., 2013). Zm00001d005082 belongs to a nodulin-like protein subfamily whose transport specificity has yet to be characterized in plants (Denancé et al., 2014). Our results suggest a potential novel role for nodulin and nodulin-like proteins, which merits further investigation.

Though rice root metaxylem were not phenotyped under drought stress, we identified several shared genetic loci that were directly or indirectly associated with drought responses. One syntenic gene pair (Zm00001d048165/Os05g0562200) encodes the protein drought-induced 19 (Table 3), which is a transcriptional activator whose downregulation mutant resulted in longer root systems and enhanced tolerance to drought and high salinity stress in *Arabidopsis* (Qin et al., 2014). Interestingly, drought-induced 19 potentially interacts with ERD15 (Qin et al., 2014), which was detected as another syntenic gene pair in the cross-species GWAS comparison. ERD15 (Zm00001d048165/Os03g0353400), is rapidly activated under drought stress and is a negative regulator of ABA (Kariola et al., 2006). ERD15 was also found to mediate osmotic stress-induced programmed cell death in soybean (*Glycine max*) protoplasts (Alves et al., 2011).

Programmed cell death elicits xylem proliferation, so it is particularly intriguing that ERD15 is associated with metaxylem vessel number in both maize and rice. Similarly, the activity of root specific kinase 1 (Zm00001d014243/Os10g0395000), which was associated with volumetric flow rate under well-watered conditions in maize and median vessel area in the *aus* rice subpopulation (Table 3), is induced by dehydration and salt (Hwang and Goodman, 1995). The maize GWAS also detected Zm00001d047710, a hypothetical gene of unknown function likely involved in ABA synthesis and degradation, that was associated with total root metaxylem vessel area under water stress (Supplemental Table S3). Though, it is surprising that more candidate genes involved in ABA metabolism were not detected in the GWAS. ABA has only recently been causally linked to induce changes in vascular development (Ramachandran et al., 2018) and crosstalk with other phytohormones creates complex interactions. More research is needed to investigate these relationships further.

Our findings suggest broadly that many genes influence root metaxylem phenotypes. In the classic paradigm, many of the candidate genes proposed here may be validated with the use of knockout or over-expression mutants to help elucidate the activity of these individual genes relative to root metaxylem phenotypes and subsequent performance outcomes under drought. However, while that strategy will certainly aid our fundamental understanding of vascular development in a crop species, it may not ultimately be helpful for producers. Our results indicate that identifying genetic loci advantageous in all environments may be difficult given the low degree of overlap between watering regimes (Supplemental Table S4). Moving forward, it may be wiser to develop new germplasm with multiplexed mutations predicted to better adapt to an unpredictable environment. Given that many root phenotypes are influenced by many genes with small effects (Schneider et al., 2020a; Schneider et al., 2020b) despite being highly heritable (Table 1), stacking multiple mutations may elicit more substantial phenotypic differences. Alternatively, we could assess the activity of our candidate genes in genotypes previously demonstrated to be more drought tolerant. Furthermore, given our limited overlap between species, a gene with demonstrated utility in one plant species may not prove to be equally as beneficial in another. Several homologs of root-specific kinase 1 (Zm00001d014243/Os10g0395000) with high sequence similarity were identified in other monocotyledonous species, but none were identified in a dicot (Figure 6). This suggests that the structure and activity of root-specific kinase 1 may have evolved to be unique to monocots and may not play a similar role in a dicotyledonous species. Therefore, we must expand our understanding of the genetic mechanisms controlling vascular development into more model systems. Vascular development specific to cereal crops has been largely overlooked and must be better characterized in order to produce novel maize lines with hydraulic systems designed to safeguard yields under stress.

In this study, root metaxylem phenotypes were observed in a panel of diverse maize lines exposed to optimal and water-limited conditions. Substantial variation was observed for all phenes measured in well-watered and water stressed conditions and these phenotypes were heritable. In a GWAS, a substantial number of gene models contained SNPs significantly associated with root metaxylem phenotypes under well-watered and water stress conditions. Few genes co-localized in both treatments, indicating that the genetic mechanisms underlying root metaxylem phenotypes are dependent on the environment. Integrating the GWAS results with a root-specific gene co-expression network facilitated prioritization of candidate genes. We argue that genes considered to be more essential in the network should be prioritized for validation. Using a cross-species GWAS comparison, we found shared genetic architecture in root metaxylem phenotypes, indicating that select genes may have conserved roles amongst species. Identifying genes that regulate root metaxylem phenotypes will deepen our understanding of plant vascular development, particularly in response to the environment, and provide novel targets for plant breeders to develop cereal varieties best suited to water-limited environments.

## Materials and Methods

### Plant Materials, Field Conditions and Experimental Design

The root phenotypes of 180 and 412 maize (*Zea mays*) genotypes from the Wisconsin Diversity Panel (Hansey et al., 2011; Hirsch et al., 2014) were evaluated in the field in 2015 and 2016, respectively, under well-watered and water stressed conditions (Supplemental Table S1). The Wisconsin Diversity Panel is a large collection of inbred maize lines of similar vigor representing the genetic diversity of North American temperate maize. A subset of 95 genotypes were grown in both 2015 and 2016. All experiments were conducted at the Apache Root Biology Center (ARBC) located near Willcox, AZ, USA (32° 29 099 N, 109° 419 3099 W). Field trials were conducted in a Grab loam soil (coarse-loamy, mixed, thermic Typic Torrifluvent) from May through October in 2015 and 2016. Each genotype was planted in a single-row plot of 20 individuals per plot with 23 cm spacing between individuals. Rows were spaced 75 cm apart (5.74 plants m^-2^). Two replications of the WiDiv were grown in each treatment in a randomized complete block design. Experiments were irrigated using a center pivot irrigation system. Well-watered and water stress treatments were induced in separate blocks. The water stress treatment was induced four weeks after planting by withholding 50% of the water needed to sustain adequate growth in the well-watered blocks. The severity of water stress was monitored over the course of the season using PR2 multi-depth soil moisture probes (Dynamax, Houston, TX, USA). Fertilizer and lime application were adjusted according to soil test results in order to meet the nutritional demands for maize production. Pest and pathogen control were applied as needed.

### Field phenotypic data

Maize root metaxylem phenotypes were evaluated using the shovelomics method (Trachsel et al., 2010) coupled with laser ablation tomography (LAT) (Hall et al., 2019; Strock et al., 2019). To summarize, the root crown of one representative maize plant was excavated from each plot just prior to anthesis in a soil monolith using a standard spade approximately 30 cm wide and deep and washed to remove the remaining soil. Samples to assess the root anatomy were collected by excising two segments from the fourth whorl nodal root 5 to 8 cm from the base of the root. The samples were preserved in 75% (v/v) ethanol in water and ablated using LAT. During LAT, the root sample is placed on an imaging stage and moved into a 355 nm pulsed laser (Avia 7000) and vaporized in the focal plane of a camera (Canon T3i with a MP-E 65mm micro lens) that simultaneously captured an image of the cross-section. Metaxylem features were quantified in each cross-section image using MIPAR software (Sosa et al., 2014). Two images were collected from each root segment and averaged to assess the mean fourth nodal root metaxylem phenotype. The theoretical volumetric flow rate was estimated based on the dimensions and number of metaxylem vessels present in each image using the Exact equations (Berker, 1963; Lewis and Boose, 1995), which estimates the volumetric movement of water through non-circular conduits.

Broad-sense heritability and repeatability on an entry mean basis of each metaxylem phene was estimated according to (Fehr, 1991). Marker-based heritability was calculated using the ‘heritability’ v.1.3 R package (Kruijer, 2019).

### Assembly of a root gene co-expression network

Co-expression analysis and module identification were conducted on a publicly available RNA-Seq dataset collected on 19 samples comprising multiple tissues and timepoints (Supplemental Table S6) (Stelpflug et al., 2016) using the WGCNA R package (Langfelder and Horvath, 2008). Genes with an FPKM < 1 for all samples were omitted from the WGCNA analysis. Parameters were adjusted so that the soft threshold was selected to produce a >90% model fit. Additionally, a minimum module size was 30 and the branch cut height was set to 0.8. Modules with a high likelihood to be related to xylem development were designated as those displaying high average gene expression in the stele (Stele_3d), differentiation zone (DifferentiationZone_3d), meristematic/elongation zone (Mz.Ez_3d), whole root system (WholeRootSystem_3d) 3 days after sowing (DAS), and zones 1 and 2 of the primary root 7 DAS (TapRoot_Z1_7d, TapRoot_Z2_7d). Gene ontology (GO) enrichment was conducted on each xylem-related module using AgriGO (Du et al., 2010) to correlate possible biological roles with xylem development.

Network topology metrics were calculated to infer the relationship between each gene and the rest of the network. For each gene, closeness centrality, i.e. the inverse of the average shortest path between the gene of interest and all other nodes in the network, and betweenness centrality, i.e. the number of shortest paths between all pairs of nodes in the network that pass through the gene of interest, were calculated using ‘igraph’ (Csardi and Nepusz, 2006). Genes in the uppermost quintile of closeness centrality and betweenness centrality were selected as hub and bottleneck genes, respectively. Genes may be labeled as both a hub and bottleneck. Network interactions were visualized using Cytoscape 3.8.0 (Lopes et al., 2010).

### Genome-wide association analysis

GWAS was performed using the phenotypic values obtained from each season and treatment and a panel of 899,784 SNPs assembled using RNA-Seq data from whole seedlings of each member of the WiDiv (Mazaheri et al., 2019) implemented in the GAPIT package (Lipka et al., 2012) in R. Single nucleotide polymorphisms (SNPs) with minor allele frequencies less than 0.01 were discarded from the panel. Due to differences in taxa membership between each season, the number of SNPs included in each GWAS differed: 578,279 in 2015 and 574,175 in 2016. The first three principal components were used as covariates to control for population structure. The minor allele effects associated with SNPs observed in the 2016 GWAS were tested using the 2015 GWAS to identify SNPs with consistent minor allele effects in both seasons. The significance threshold was selected using a linear model that predicts a significant p-threshold according to the marker-based heritability (Kaler and Purcell, 2019). Using the mean marker-based heritability of all phenes (0.264), the p-threshold was predicted to be at -log(p) ≤ 3.43. A SNP that satisfied the p-threshold in 2016 and displayed consistent minor allele effects in both seasons was designated as “significant.” Candidate genes corresponding to a significant SNP were detected according to the AGPv4 reference sequence assembly (Jiao et al., 2017). Annotation of candidate genes was obtained from MaizeGDB (Portwood et al., 2018).

### Comparative GWAS between Maize and Rice

GWAS was conducted on rice metaxylem phenes grown under well-watered conditions in the greenhouse (Reeger et al., unpublished). Rice root anatomy was evaluated on 336 rice (*Oryza sativa*) accessions from the Rice Diversity Panel 1 (RDP1) (Zhao et al., 2011) grown in greenhouses at University Park, PA (40°49’ N, 77°52’ W). Accessions represented all *O. sativa* subpopulations: 52 *aus*, 67 *indica*, 11 *aromatic*, 74 *temperate japonica*, 80 *tropical japonica*, and 51 admixed accessions. Plants were grown in 10.5 L pots (21 cm x 40.6 cm, top diameter x height, Nursery Supplies Inc., Chambersburg, PA, USA) filled with growth medium consisting of 40% v/v medium size (0.3-0.5 mm) commercial grade sand (Quikrete Companies Inc., Harrisburg, PA, USA), 60% v/v horticultural vermiculite (Whittemore Companies Inc., Lawrence, MA, USA), and solid-phase buffered phosphorus (Al-P, prepared according to (Lynch et al., 1990)) providing a constant availability of high (100 μM) phosphorus concentration in the soil solution. Each pot was direct-seeded with three seeds, and after 7 days plants were thinned to one per pot. Plants were irrigated once per day via drip irrigation with Yoshida nutrient solution without phosphate (Yoshida et al., 1971), which was adjusted to pH 5.5-6.0 daily. At the 8th leaf stage, root systems of each plant were excavated, washed with water, and preserved in 70% (v/v) ethanol in water for later sampling. Preserved root tissue at 15 cm from the root apex of two nodal roots was freehand sectioned under a dissection microscope (SMZ-U, Nikon, Tokyo, Japan) at 4x magnification using Teflon-coated double-edged stainless steel blades (Electron Microscopy Sciences, Hatfield, PA, USA). Images of the root cross-section were captured using a Diaphot inverted light microscope (SMZ-U, Nikon, Tokyo, Japan) at 40x magnification equipped with digital camera (NIKON DS-Fi1, Tokyo, Japan). The two best images were selected to quantify root anatomy using the image analysis software Rice Root Analyzer (Taeparsartsit 2013, unpublished) to measure the number of metaxylem vessels and RootScan software (Burton et al., 2012) to measure metaxylem vessel area. Mean values of each root anatomical phene for each genotype were calculated and used for all further analysis.

Using the 4.8 million SNPs from the Rice Reference Panel (RICE-RP) (Wang et al., 2018), genome-wide associations were calculated for root anatomical traits in all individuals (ALL), as well as within subpopulations (*indica*, *IND*; *tropical-japonica*, *TRJ*; *temperate-japonica*, *TEJ*; aus, *AUS*). To avoid false-positives and reduce computational requirements per association, the SNPs were divided into 6 non-orthogonal subsets by linkage-disequilibrium (r^2^) pruning using the *indep* and *indep_pairwise* functions in plink1.9 (Chang et al., 2015). A linear mixed-model in the gwas() function of the ‘rrBLUP’ package in R was used for all associations (Endelman, 2011). A minimum minor-allele frequency of 0.05 was used (min.MAF = 0.05) with at least 3 minor allele count, and variance components were calculated once (P3D = TRUE). Based on previous methods (McCouch et al., 2016), no principle components were used for single subpopulations, but three principle components were added for associations in ALL (n.PC = 3). To identify genomic regions in the output, the following requirements were used: region had greater than 3 SNPs with -log(*p*) > 4 that were within 200 kb of each other, region was significant in at least 5 of 6 SNP subsets, and the most-significant SNP (MS-SNP) in the region had -log(*p*) > 5. In total, 3636 candidate genes were detected in rice for median metaxylem area and number of metaxylem vessels. Genes within these regions with their GO names were retrieved using Bioconductor tools in R (Huber et al., 2015).

Maize and rice syntenic genes were identified using the EnsemblPlants database of homologs via the biomaRt package in R (Huber et al., 2015), which utilizes a synteny method based on orthology that was previously described (Schnable et al., 2009). Syntenic gene pairs most likely associated with root metaxylem phenes were made up of genes with a significant association in both maize (-log(*p*) > 3.43) and rice (-log(*p*) > 4) GWAS. Significant allelic effects on maize root metaxylem phenotypes were determined using a two-sample t-test (α < 0.05). Because rice candidate genes may be distant from the significant SNP associated with a root metaxylem phene, haplotypes of each candidate gene were identified using a cluster analysis and their metaxylem root phenotypes were compared. SNP genotypes across accessions, which included all nonsynonymous SNPs in the candidate gene as determined by the SNP-Seek database (Mansueto et al., 2017), were clustered into similar groups using K-means clustering via the ‘pheatmap’ v1.0.12 package in R (Kolde, 2015), where the number of clusters was determined using the within sums of squares method. Significant differences in rice root metaxylem phenotypes across haplotypes were determined using Kruskal-Wallis and pairwise Wilcoxon multiple comparison tests (α < 0.05). Phylogenetic trees based on syntenic gene pairs were constructed to infer the evolutionary history. Sequences of high similarity (megablast) to the maize gene were identified using BLAST and their amino acid sequences were aligned using the MUSCLE algorithm in MEGA X (Hall, 2013). All positions with less than 95% site coverage were eliminated. A bootstrap consensus tree was constructed from 500 replications of Maximum Likelihood trees in MEGA X (Hall, 2013) and edited in TreeGraph 2 (Stöver and Müller, 2010). The percentage of replicate trees in which the associated taxa clustered together is shown next to the branches.

### Data Analysis

All data analyses were performed in R version 4.0.0 (R Core Team, 2020) with graphical illustration using ‘ggplot2’ (Wickham, 2016).

## Supplemental Data

Supplemental Table S1. Genotypes of the Wisconsin Diversity Panel grown in 2015 and 2016.

Supplemental Table S2. Number of SNPs exhibiting consistent effects on root phenotypes in each treatment across both seasons.

Supplemental Table S3. Significant SNPs and candidate genes identified via GWAS for associated with all root metaxylem phenes under well-watered and water stress conditions. MXAmax, maximum root metaxylem vessel area; MXAmed, median root metaxylem vessel area; MXAtot, total root metaxylem vessel area; MXN, metaxylem vessel number; VFR, volumetric flow rate.

Supplemental Table S4. Significant SNPs and candidate genes that were associated with root metaxylem phenes in both well-watered (WW) and water stress (WS) conditions. MXAmax, maximum root metaxylem vessel area; MXAmed, median root metaxylem vessel area; MXAtot, total root metaxylem vessel area; MXN, metaxylem vessel number; VFR, volumetric flow rate.

Supplemental Table S5. Statistics describing network topology and membership size for each co-expression module strongly associated with root tissues relevant to root metaxylem development.

Supplemental Table S6. List of 19 root tissues used to build the root-specific gene co-expression network.

Supplemental Figure S1. Correlations between mean values of each genotype under well-watered (WW) and water stressed (WS) conditions for each metaxylem phene in 2015 (black) and 2016 (grey). The solid black or grey lines indicate the slope of the correlation. The solid blue line indicates where x = y.

Supplemental Figure S2. Q-Q plots assessing the fitness of the model for GWAS of root metaxylem phenes under well-watered (WW) and water stress (WS) conditions.

Supplemental Figure S3. General outline of the filtering steps used to identify SNPs with consistent allelic effects in separate GWAS conducted with data collected in 2015 and 2015. (A) A substantial proportion of the SNP panel was included in each GWAS. Binned scatterplots showing (B) the full SNP panel and (C) the filtered set containing SNPs of consistent minor allele effects. The gradation of the bin corresponds to the number of SNPs residing in the bin where densely populated bins are light blue while more sparsely populated bins are dark blue.

Supplemental Figure S4. Mapman ontogenic categories for annotated gene models associated with significant SNPs in well-watered and water stress conditions.

Supplemental Figure S5. Network topology metrics used to identify hub and bottleneck genes in the pool of candidate genes identified via GWAS. Scatterplots showing closeness centrality against betweenness centrality of each gene for the remaining gene co-expression modules most strongly associated with root tissues relevant to metaxylem development. Modules “navajowhite2” and “thistle1” did not contain any significant SNPs detected in the GWAS. Colors correspond to bottleneck (BN, red), bottleneck/hub (BN/HUB, yellow), hub (HUB, blue), and non-affiliated (None, grey) genes.

Supplemental Figure S6. Haplotypes of rice genes encoding a root-specific kinase 1 (Os10g0395000).

## Acknowledgements

This research was supported by the National Institute of Food and Agriculture, U.S. Department of Agriculture, award #2017-67013-26192 and Hatch project 4732, and the Howard G. Buffett Foundation. We thank Robert Snyder and Patricio Cid for technical support; and Charles Anderson, Ivan Baxter Brian Dilkes, and Jesse Lasky for helpful discussions.

## Author contributions

SPK designed the experiments, analyzed the data, and wrote the article with contributions from all authors; JER analyzed data and contributed to writing; SMK and KMB provided feedback on data analysis and writing; JPL conceived and supervised the project and contributed to data analysis and writing. JPL agrees to serve as the author responsible for contact and ensures communication.

## Abbreviations

ABA: abscisic acid
BN: bottleneck gene
BN/HUB: bottleneck and hub gene
GWAS: genome-wide association study
HUB: hub gene
SNP: single nucleotide polymorphism
WS: water stress
WW: well-watered

## References

Abd Allah AA, Badawy SA, Zayed BA, El. Gohary AA (2010) The Role of Root System Traits in the Drought Tolerance of Rice (Oryza sativa L.). International Journal of Agricultural and Biological Sciences 1: 83–87

Alves MS, Reis PAB, Dadalto SP, Faria JAQA, Fontes EPB, Fietto LG (2011) A novel transcription factor, ERD15 (Early Responsive to Dehydration 15), connects endoplasmic reticulum stress with an osmotic stress-induced cell death signal. J Biol Chem 286: 20020–20030

Balestrazzi A, Confalonieri M, Macovei A, Donà M, Carbonera D (2011) Genotoxic stress and DNA repair in plants: emerging functions and tools for improving crop productivity. Plant Cell Rep 30: 287–295

Bekh-Ochir D, Shimada S, Yamagami A, Kanda S, Ogawa K, Nakazawa M, Matsui M, Sakuta M, Osada H, Asami T, et al (2013) A novel mitochondrial DnaJ/Hsp40 family protein BIL2 promotes plant growth and resistance against environmental stress in brassinosteroid signaling. Planta 237: 1509–1525

Bennetzen JL, Freeling M (1997) The unified grass genome: synergy in synteny. Genome Res 7:301–306

Berker R (1963) Intégration des équations du mouvement d’un fluide visqueux incompressible. Handbuch der Physik 3: 1–384

Buell CR, Yuan Q, Ouyang S, Liu J, Zhu W, Wang A, Maiti R, Haas B, Wortman J, Pertea M, et al (2005) Sequence, annotation, and analysis of synteny between rice chromosome 3 and diverged grass species. Genome Res 15: 1284–1291

Burridge JD, Rangarajan H, Lynch JP (2020) Comparative phenomics of annual grain legume root architecture. Crop Sci 1–20

Burton AL, Williams M, Lynch JP, Brown KM (2012) RootScan: Software for high-throughput analysis of root anatomical traits. Plant Soil 357: 189–203

Carpita NC (1996) Structure and biogenesis of the cell walls of grasses. Annual Reviews in Plant Physiology Plant Molecular Biology 47: 445–476

Chang CC, Chow CC, Tellier LC, Vattikuti S, Purcell SM, Lee JJ (2015) Second-generation PLINK: rising to the challenge of larger and richer datasets. Gigascience 4: 1 –16

Chen W, Wang W, Peng M, Gong L, Gao Y, Wan J, Wang S, Shi L, Zhou B, Li Z, et al (2016) Comparative and parallel genome-wide association studies for metabolic and agronomic traits in cereals. Nature Communciations 7: 1 –10

Chimungu JG, Brown KM, Lynch JP (2014a) Large root cortical cell size improves drought tolerance in maize. Plant Physiol 166: 2166–2178

Chimungu JG, Brown KM, Lynch JP (2014b) Reduced root cortical cell file number improves drought tolerance in maize. Plant Physiol 166: 1943–1955

Choe S, Dilkes BP, Gregory BD, Ross AS, Yuan H, Noguchi T, Fujioka S, Takatsuto S, Tanaka A, Yoshida S, et al (1999a) The Arabidopsis dwarf1 mutant is defective in the conversion of 24-methylenecholesterol to campesterol in brassinosteroid biosynthesis. Plant Physiol 119: 897–907

Choe S, Noguchi T, Fujioka S, Takatsuto S, Tissier CP, Gregory BD, Ross AS, Tanaka A, Yoshida S, Tax FE, et al (1999b) The Arabidopsis dwf7/ste1 mutant is defective in the delta7 sterol C-5 desaturation step leading to brassinosteroid biosynthesis. Plant Cell 11: 207–221

Comas LH, Becker SR, Cruz VMV, Byrne PF, Dierig DA (2013) Root traits contributing to plant productivity under drought. Front Plant Sci 4: 1–16

Csardi G, Nepusz T (2006) The igraph software package for complex network research. InterJournal, Complex Systems 1695

Dai A (2013) Increasing drought under global warming in observations and models. Nat Clim Chang 3: 52–58

Daryanto S, Wang L, Jacinthe P-A (2016) Global Synthesis of Drought Effects on Maize and Wheat Production. PLoS One 11: e0156362

Denancé N, Szurek B, Noël LD (2014) Emerging functions of nodulin-like proteins in non-nodulating plant species. Plant Cell Physiol 55: 469–474

Du Z, Zhou X, Ling Y, Zhang Z, Su Z (2010) agriGO: a GO analysis toolkit for the agricultural community. Nucleic Acids Res 38: W64–70

Endelman JB (2011) Ridge regression and other kernels for genomic selection with R package rrBLUP. Plant Genome 4: 250–255

Fehr W (1991) Principles of Cultivar Development: Theory and Technique. Iowa State University

Feng W, Lindner H, Robbins NE 2nd, Dinneny JR (2016) Growing Out of Stress: The Role of Cell- and Organ-Scale Growth Control in Plant Water-Stress Responses. Plant Cell 28: 1769–1782

Finkelstein R (2013) Abscisic Acid synthesis and response. *In* C Somerville, E Meyerowitz, eds, The Arabidopsis Book. pp 1–36

Gao Y, Lynch JP (2016) Reduced crown root number improves water acquisition under water deficit stress in maize (Zea mays L.). J Exp Bot 67: 4545–4557

Guet J, Fichot R, Lédée C, Laurans F, Cochard H, Delzon S, Bastien C, Brignolas F (2015) Stem xylem resistance to cavitation is related to xylem structure but not to growth and water-use efficiency at the within-population level in Populus nigra L. J Exp Bot 66: 4643–4652

Guo H, Li L, Ye H, Yu X, Algreen A, Yin Y (2009a) Three related receptor-like kinases are required for optimal cell elongation in Arabidopsis thaliana. Proc Natl Acad Sci U S A 106: 7648–7653

Guo H, Ye H, Li L, Yin Y (2009b) A family of receptor-like kinases are regulated by BES1 and involved in plant growth in Arabidopsis thaliana. Plant Signal Behav 4: 784–786

Hacke UG, Sperry JS (2001) Functional and ecological xylem anatomy. Perspect Plant Ecol Evol Syst 4: 97–115

Hall BG (2013) Building phylogenetic trees from molecular data with MEGA. Mol Biol Evol 30:1229–1235

Hall B, Lanba A, Lynch J (2019) Three-dimensional analysis of biological systems via a novel laser ablation technique. J Laser Appl 31: 022602

Hansey CH, Johnson JM, Sekhon RS, Kaeppler SM, de Leon N (2011) Genetic diversity of a maize association population with restricted phenology. Crop Sci 51: 704–715

Henry A, Cal AJ, Batoto TC, Torres RO, Serraj R (2012) Root attributes affecting water uptake of rice (Oryza sativa) under drought. J Exp Bot 63: 4751–4763

Hirsch CN, Foerster JM, Johnson JM, Sekhon RS, Muttoni G, Vaillancourt B, Peñagaricano F, Lindquist E, Pedraza MA, Barry K, et al (2014) Insights into the maize pan-genome and pan-transcriptome. Plant Cell 26: 121–135

Hochholdinger F (2009) The Maize Root System: Morphology, Anatomy, and Genetics. Handbook of Maize: Its Biology. Springer New York, pp 145–160

Ho MD, Rosas JC, Brown KM, Lynch JP (2005) Root architectural tradeoffs for water and phosphorus acquisition. Funct Plant Biol 32: 737–748

Hose E, Steudle E, Hartung W (2000) Abscisic acid and hydraulic conductivity of maize roots: a study using cell- and root-pressure probes. Planta 211: 874–882

Huang J, Kim CM, Xuan Y-H, Park SJ, Piao HL, Je BI, Liu J, Kim TH, Kim B-K, Han C-D (2013) OsSNDP1, a Sec14-nodulin domain-containing protein, plays a critical role in root hair elongation in rice. Plant Mol Biol 82: 39–50

Huber W, Carey VJ, Gentleman R, Anders S, Carlson M, Carvalho BS, Bravo HC, Davis S, Gatto L, Girke T, et al (2015) Orchestrating high-throughput genomic analysis with Bioconductor. Nat Methods 12: 115–121

Ingvarsson PK, Street NR (2011) Association genetics of complex traits in plants: Tansley review. New Phytol 189: 909–922

IPCC (2013) Climate Change 2013: The Physical Science Basis. Contribution of Working Group I to the Fifth Assessment Report of the Intergovernmental Panel on Climate Change. Cambridge University Press

Jaramillo RE, Nord EA, Chimungu JG, Brown KM, Lynch JP (2013) Root cortical burden influences drought tolerance in maize. Ann Bot 112: 429–437

Jeong H, Mason SP, Barabási A-L, Oltvai ZN (2001) Lethality and centrality in protein networks. Nature 411: 41–42

Jiao Y, Peluso P, Shi J, Liang T, Stitzer MC, Wang B, Campbell MS, Stein JC, Wei X, Chin C-S, et al (2017) Improved maize reference genome with single-molecule technologies. Nature 546: 524–527

Johnsson C, Jin X, Xue W, Dubreuil C, Lezhneva L, Fischer U (2019) The plant hormone auxin directs timing of xylem development by inhibition of secondary cell wall deposition through repression of secondary wall NAC-domain transcription factors. Physiol Plant 165: 673–689

Kadam NN, Tamilselvan A, Lawas LMF, Quinones C, Bahuguna RN, Thomson MJ, Dingkuhn M, Muthurajan R, Struik PC, Yin X, et al (2017) Genetic Control of Plasticity in Root Morphology and Anatomy of Rice in Response to Water Deficit. Plant Physiol 174: 2302–2315

Kadam NN, Yin X, Bindraban PS, Struik PC, Jagadish KSV (2015) Does morphological and anatomical plasticity during the vegetative stage make wheat more tolerant of water deficit stress than rice? Plant Physiol 167: 1389–1401

Kaler AS, Purcell LC (2019) Estimation of a significance threshold for genome-wide associated studies. BMC Genomics 20: 1 –8

Kano M, Inukai Y, Kitano H, Yamauchi A (2011) Root plasticity as the key root trait for adaptation to various intensities of drought stress in rice. Plant Soil 342: 117–128

Kariola T, Brader G, Helenius E, Li J, Heino P, Palva ET (2006) EARLY RESPONSIVE TO DEHYDRATION 15, a negative regulator of abscisic acid responses in Arabidopsis. Plant Physiol 142: 1559–1573

Klein SP, Schneider HM, Perkins AC, Brown KM, Lynch JP (2020) Multiple Integrated Root Phenotypes Are Associated with Improved Drought Tolerance. Plant Physiol 183: 1011 –1025

Kolde R (2015) pheatmap: Pretty heatmaps.

Kruijer W (2019) heritability: Marker-Based Estimation of Heritability Using Individual Plant or Plot Data.

Kubo M, Udagawa M, Nishikubo N, Horiguchi G, Yamaguchi M, Ito J, Mimura T, Fukuda H, Demura T (2005) Transcription switches for protoxylem and metaxylem vessel formation. Genes Dev 19: 1855–1860

Langfelder P, Horvath S (2008) WGCNA: an R package for weighted correlation network analysis. BMC Bioinformatics 9: 1 –13

Lewis AM, Boose ER (1995) Estimating Volume Flow Rates Through Xylem Conduits. Am J Bot 82: 1112–1116

Liakat Ali M, Luetchens J, Nascimento J, Shaver TM, Kruger GR, Lorenz AJ (2015) Genetic variation in seminal and nodal root angle and their association with grain yield of maize under water-stressed field conditions. Plant and Soil 397: 213–225

Li H, Zhang D, Wang X, Li H, Rengel Z, Shen J (2018) Competition between Zea mays genotypes with different root morphological and physiological traits is dependent on phosphorus forms and supply patterns. Plant Soil 434: 125–137

Lipka AE, Tian F, Wang Q, Peiffer J, Li M, Bradbury PJ, Gore MA, Buckler ES, Zhang Z (2012) GAPIT: genome association and prediction integrated tool. Bioinformatics 28: 2397–2399

Lobell DB, Roberts MJ, Schlenker W, Braun N, Little BB, Rejesus RM, Hammer GL (2014) Greater sensitivity to drought accompanies maize yield increase in the U.S. Midwest. Science 344: 516–519

Lobell DB, Schlenker W, Costa-Roberts J (2011) Climate trends and global crop production since 1980. Science 333: 616–620

Lopes CT, Franz M, Kazi F, Donaldson SL, Morris Q, Bader GD (2010) Cytoscape Web: an interactive web-based network browser. Bioinformatics 26: 2347–2348

Lynch J, Epstein E, Lauchli A, Weight GI (1990) An automated greenhouse sand culture system suitable for studies of P nutrition. Plant Cell Environ 13: 547–554

Lynch JP (2013) Steep, cheap and deep: an ideotype to optimize water and N acquisition by maize root systems. Ann Bot 112: 347–357

Lynch JP (2019) Root phenotypes for improved nutrient capture: an underexploited opportunity for global agriculture. New Phytol 223: 548–564

Lynch JP (2018) Rightsizing root phenotypes for drought resistance. J Exp Bot 69: 3279–3292

Lynch JP, Chimungu JG, Brown KM (2014) Root anatomical phenes associated with water acquisition from drying soil: targets for crop improvement. J Exp Bot 65: 6155–6166

Mansueto L, Fuentes RR, Borja FN, Detras J, Abriol-Santos JM, Chebotarov D, Sanciangco M, Palis K, Copetti D, Poliakov A, et al (2017) Rice SNP-seek database update: new SNPs, indels, and queries. Nucleic Acids Res 45: D1075–D1081

Maurel C, Nacry P (2020) Root architecture and hydraulics converge for acclimation to changing water availability. Nat Plants 6: 744–749

Mazaheri M, Heckwolf M, Vaillancourt B, Gage JL, Burdo B, Heckwolf S, Barry K, Lipzen A, Ribeiro CB, Kono TJY, et al (2019) Genome-wide association analysis of stalk biomass and anatomical traits in maize. BMC Plant Biology 19: 1–17

McCouch SR, Wright MH, Tung C-W, Maron LG, McNally KL, Fitzgerald M, Singh N, DeClerck G, Agosto-Perez F, Korniliev P, et al (2016) Open access resources for genomewide association mapping in rice. Nat Commun 7: 10532

McDermott JE, Taylor RC, Yoon H, Heffron F (2009) Bottlenecks and hubs in inferred networks are important for virulence in Salmonella typhimurium. J Comput Biol 16: 169–180

McLaren W, Gil L, Hunt SE, Riat HS, Ritchie GRS, Thormann A, Flicek P, Cunningham F (2016) The Ensembl Variant Effect Predictor. Genome Biol 17: 1–14

Milhinhos A, Miguel CM (2013) Hormone interactions in xylem development: a matter of signals. Plant Cell Reports 32: 867–883

Okekeogbu IO, Pattathil S, González Fernández-Niño SM, Aryal UK, Penning BW, Lao J, Heazlewood JL, Hahn MG, McCann MC, Carpita NC (2019) Glycome and Proteome Components of Golgi Membranes Are Common between Two Angiosperms with Distinct Cell-Wall Structures. Plant Cell 31: 1094–1112

Oyiga BC, Palczak J, Wojciechowski T, Lynch JP, Naz AA, Léon J, Ballvora A (2020) Genetic components of root architecture and anatomy adjustments to water-deficit stress in spring barley. Plant Cell Environ 43: 692–711

Parent B, Hachez C, Redondo E, Simonneau T, Chaumont F, Tardieu F (2009) Drought and abscisic acid effects on aquaporin content translate into changes in hydraulic conductivity and leaf growth rate: a trans-scale approach. Plant Physiol 149: 2000–2012

Pesquet E, Tuominen H (2011) Ethylene stimulates tracheary element differentiation in Zinnia elegans cell cultures. New Phytol 190: 138–149

Pires MV, de Castro EM, de Freitas BSM, Souza Lira JM, Magalhães PC, Pereira MP (2020) Yield-related phenotypic traits of drought resistant maize genotypes. Environ Exp Bot 171: 1–10

Portwood JL, Woodhouse MR, Cannon EK, Gardiner JM, Harper LC, Schaeffer ML, Walsh JR, Sen TZ, Cho KT, Schott DA, et al (2018) MaizeGDB 2018: the maize multi-genome genetics and genomics database. Nucleic Acids Research 47: 1146–1154

Purushothaman R, Zaman-Allah M, Mallikarjuna N, Pannirselvam R, Krishnamurthy L, Gowda CLL (2013) Root Anatomical Traits and Their Possible Contribution to Drought Tolerance in Grain Legumes. Plant Prod Sci 16: 1–8

Qin L-X, Li Y, Li D-D, Xu W-L, Zheng Y, Li X-B (2014) Arabidopsis drought-induced protein Di19-3 participates in plant response to drought and high salinity stresses. Plant Mol Biol 86: 609–625

Ramachandran P, Augstein F, Nguyen V, Carlsbecker A (2020) Coping With Water Limitation: Hormones That Modify Plant Root Xylem Development. Front Plant Sci 11: 570

Ramachandran P, Wang G, Augstein F, de Vries J, Carlsbecker A (2018) Continuous root xylem formation and vascular acclimation to water deficit involves endodermal ABA signalling via miR165. Development 145: 1-7

R Core Team (2020) R: A language and environment for statistical computing. R Foundation for Statistical Computing, Vienna, Austria

Richards RA, Passioura JB (1989) A breeding program to reduce the diameter of the major xylem vessel in the seminal roots of wheat and its effect on grain-yield in rain-fed environments. Aust J Agric Res 40: 943–950

Richmond TA, Somerville CR (2000) The cellulose synthase superfamily. Plant Physiol 124: 495–498

Roberts K, McCann MC (2000) Xylogenesis: the birth of a corpse. Curr Opin Plant Biol 3: 517–522

Růžička K, Ursache R, Hejátko J, Helariutta Y (2015) Xylem development - from the cradle to the grave. New Phytol 207: 519–535

Salse J, Piegu B, Cooke R, Delseny M (2004) New in silico insight into the synteny between rice (Oryza sativa L.) and maize (Zea mays L.) highlights reshuffling and identifies new duplications in the rice genome. The Plant Journal 38: 396–409

Schaefer RJ, Michno J-M, Jeffers J, Hoekenga O, Dilkes B, Baxter I, Myers CL (2018) Integrating Coexpression Networks with GWAS to Prioritize Causal Genes in Maize. Plant Cell 30: 2922–2942

Schaefer RJ, Michno J-M, Myers CL (2017) Unraveling gene function in agricultural species using gene co-expression networks. Biochim Biophys Acta Gene Regul Mech 1860: 53–63

Schnable PS, Ware D, Fulton RS, Stein JC, Wei F, Pasternak S, Liang C, Zhang J, Fulton L, Graves TA, et al (2009) The B73 Maize Genome: Complexity, Diversity, and Dynamics. Science 326: 1112–1115

Schneider HM, Klein SP, Hanlon MT, Kaeppler S, Brown KM, Lynch JP (2020a) Genetic control of root anatomical plasticity in maize. Plant Genome 13: 1–14

Schneider HM, Klein SP, Hanlon MT, Nord EA, Kaeppler S, Brown KM, Warry A, Bhosale R, Lynch JP (2020b) Genetic Control of Root Architectural Plasticity in Maize. J Exp Bot 71: 3185–3197

Schneider HM, Lynch JP (2020) Should Root Plasticity Be a Crop Breeding Target? Front Plant Sci 11: 1–16

Schuetz M, Smith R, Ellis B (2012) Xylem tissue specification, patterning, and differentiation mechanisms. J Exp Bot 64: 11–31

Smith BG, Harris PJ (1999) The polysaccharide composition of Poales cell walls: Poaceae cell walls are not unique. Biochem Syst Ecol 27: 33–53

Sosa JM, Huber DE, Welk B, Fraser HL (2014) Development and application of MIPAR™: a novel software package for two- and three-dimensional microstructural characterization. Integrating Materials and Manufacturing Innovation 3: 123–140

de Souza TC, de Castro EM, César Magalhães P, De Oliveira Lino L, Trindade Alves E, de Albuquerque PEP (2013) Morphophysiology, morphoanatomy, and grain yield under field conditions for two maize hybrids with contrasting response to drought stress. Acta Physiol Plant 35: 3201–3211

Sperry JS, Saliendra NZ (1994) Intra- and inter-plant variation in xylem cavitation in Betula occidentalis. Plant Cell Environ 17: 1233–1241

Stelpflug SC, Sekhon RS, Vaillancourt B, Hirsch CN, Buell CR, de Leon N, Kaeppler SM (2016) An Expanded Maize Gene Expression Atlas based on RNA Sequencing and its Use to Explore Root Development. Plant Genome 9: 1 –16

Stöver BC, Müller KF (2010) TreeGraph 2: Combining and visualizing evidence from different phylogenetic analyses. BMC Bioinformatics 11: 1–9

Strock CF, Schneider HM, Galindo-Castañeda T, Hall BT, Van Gansbeke B, Mather DE, Roth MG, Chilvers MI, Guo X, Brown K, et al (2019) Laser ablation tomography for visualization of root colonization by edaphic organisms. Journal of Experimental Botany 70: 5327–5342

Szekeres M, Németh K, Koncz-Kálmán Z, Mathur J, Kauschmann A, Altmann T, Rédei GP, Nagy F, Schell J, Koncz C (1996) Brassinosteroids rescue the deficiency of CYP90, a cytochrome P450, controlling cell elongation and de-etiolation in Arabidopsis. Cell 85: 171–182

Taylor NG, Howells RM, Huttly AK, Vickers K, Turner SR (2003) Interactions among three distinct CesA proteins essential for cellulose synthesis. Proc Natl Acad Sci U S A 100: 1450–1455

Trachsel S, Kaeppler SM, Brown KM, Lynch JP (2010) Shovelomics: high throughput phenotyping of maize (Zea mays L.) root architecture in the field. Plant Soil 341: 75–87

Turco GM, Rodriguez-Medina J, Siebert S, Han D, Valderrama-Gómez MÁ, Vahldick H, Shulse CN, Cole BJ, Juliano CE, Dickel DE, et al (2019) Molecular Mechanisms Driving Switch Behavior in Xylem Cell Differentiation. Cell Rep 28: 342–351.e4

Uga Y, Sugimoto K, Ogawa S, Rane J, Ishitani M, Hara N, Kitomi Y, Inukai Y, Ono K, Kanno N, et al (2013) Control of root system architecture by DEEPER ROOTING 1 increases rice yield under drought conditions. Nat Genet 45: 1097–1102

Vadez V (2014) Root hydraulics: The forgotten side of roots in drought adaptation. Field Crops Res 165:15–24

Vadez V, Kholova J, Medina S, Kakkera A, Anderberg H (2014) Transpiration efficiency: new insights into an old story. J Exp Bot 65: 6141–6153

Vejchasarn P (2014) Nutritional and Genetic Architecture of Root Traits in Rice (Oryza sativa). Ph.D. The Pennsylvania State University.

Vilagrosa A, Chirino E, Peguero-Pina JJ, Barigah TS, Cochard H, Gil-Pelegrín E (2012) Xylem Cavitation and Embolism in Plants Living in Water-Limited Ecosystems. *In* R Aroca, ed, Plant Responses to Drought. Springer-Verlag, pp 63–109

Vogel J (2008) Unique aspects of the grass cell wall. Curr Opin Plant Biol 11: 301–307

Wahl S, Ryser P (2000) Root tissue structure is linked to ecological strategies of grasses. New Phytol 148: 459–471

Wang DR, Agosto-Pérez FJ, Chebotarov D, Shi Y, Marchini J, Fitzgerald M, McNally KL, Alexandrov N, McCouch SR (2018) An imputation platform to enhance integration of rice genetic resources. Nat Commun 9: 3519

Wasson AP, Richards RA, Chatrath R, Misra SC, Prasad SVS, Rebetzke GJ, Kirkegaard JA, Christopher J, Watt M (2012) Traits and selection strategies to improve root systems and water uptake in water-limited wheat crops. J Exp Bot 63: 3485–3498

Wickham H (2016) ggplot2: Elegant Graphics for Data Analysis. Springer-Verlag, New York

Yamaguchi M, Goué N, Igarashi H, Ohtani M, Nakano Y, Mortimer JC, Nishikubo N, Kubo M, Katayama Y, Kakegawa K, et al (2010a) VASCULAR-RELATED NAC-DOMAIN6 and VASCULAR-RELATED NAC-DOMAIN7 effectively induce transdifferentiation into xylem vessel elements under control of an induction system. Plant Physiol 153: 906–914

Yamaguchi M, Mitsuda N, Ohtani M, Ohme-Takagi M, Kato K, Demura T (2011) VASCULAR-RELATED NAC-DOMAIN 7 directly regulates the expression of a broad range of genes for xylem vessel formation: Direct target genes of VND7. Plant J 66: 579–590

Yamaguchi M, Ohtani M, Mitsuda N, Kubo M, Ohme-Takagi M, Fukuda H, Demura T (2010b) VND-INTERACTING2, a NAC domain transcription factor, negatively regulates xylem vessel formation in Arabidopsis. Plant Cell 22: 1249–1263

Yoshida S, Forno DA, Bhadrachalam A (1971) Zinc deficiency of the rice plant on calcareous and neutral soils in the philippines. Soil Sci Plant Nutr 17: 83–87

Yu H, Greenbaum D, Xin Lu H, Zhu X, Gerstein M (2004) Genomic analysis of essentiality within protein networks. Trends Genet 20: 227–231

Yu H, Kim PM, Sprecher E, Trifonov V, Gerstein M (2007) The importance of bottlenecks in protein networks: correlation with gene essentiality and expression dynamics. PLoS Comput Biol 3: e59

Zaman-Allah M, Jenkinson DM, Vadez V (2011) A conservative pattern of water use, rather than deep or profuse rooting, is critical for the terminal drought tolerance of chickpea. J Exp Bot 62: 4239–4252

Zhan A, Schneider H, Lynch JP (2015) Reduced Lateral Root Branching Density Improves Drought Tolerance in Maize. Plant Physiol 168: 1603–1615

Zhang X, Pang J, Ma X, Zhang Z, He Y, Hirsch CN, Zhao J (2019) Multivariate analyses of root phenotype and dynamic transcriptome underscore valuable root traits and water-deficit responsive gene networks in maize. Plant Direct 3: 1–18

Zhao K, Tung C-W, Eizenga GC, Wright MH, Ali ML, Price AH, Norton GJ, Islam MR, Reynolds A, Mezey J, et al (2011) Genome-wide association mapping reveals a rich genetic architecture of complex traits in Oryza sativa. Nat Commun 2: 467

Zheng Z, Hey S, Jubery T, Liu H, Yang Y, Coffey L, Miao C, Sigmon B, Schnable JC, Hochholdinger F, et al (2020) Shared Genetic Control of Root System Architecture between Zea mays and Sorghum bicolor. Plant Physiol 182: 977–991

Zhu J, Brown KM, Lynch JP (2010) Root cortical aerenchyma improves the drought tolerance of maize (Zea mays L.). Plant Cell Environ 9: 31

