## Supplemental Figures for "Shared genetic architecture underlying root metaxylem phenotypes under drought stress in cereals"

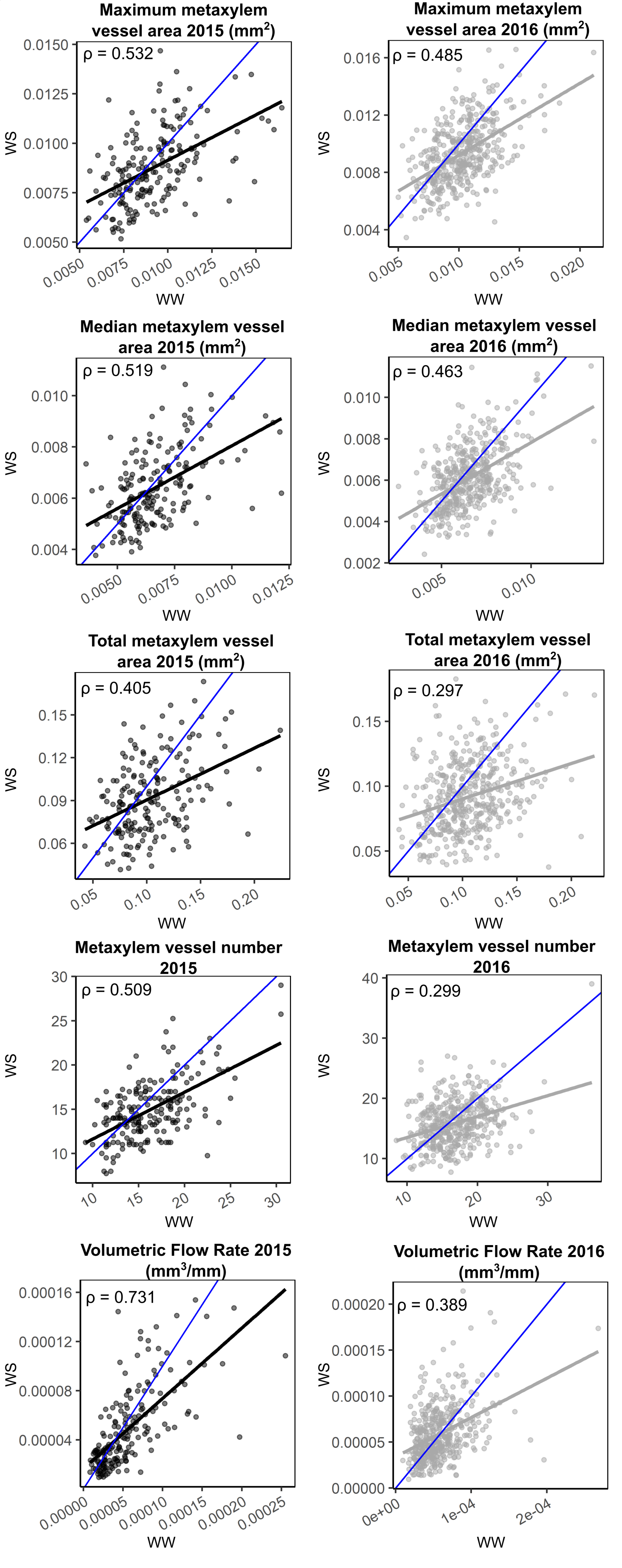


Supplemental Figure S1. Correlations between mean values of each genotype under well-watered (WW) and water stressed (WS) conditions for each metaxylem phene in 2015 (black) and 2016 (grey). The solid black or grey lines indicate the slope of the correlation. The solid blue line indicates where x = y.

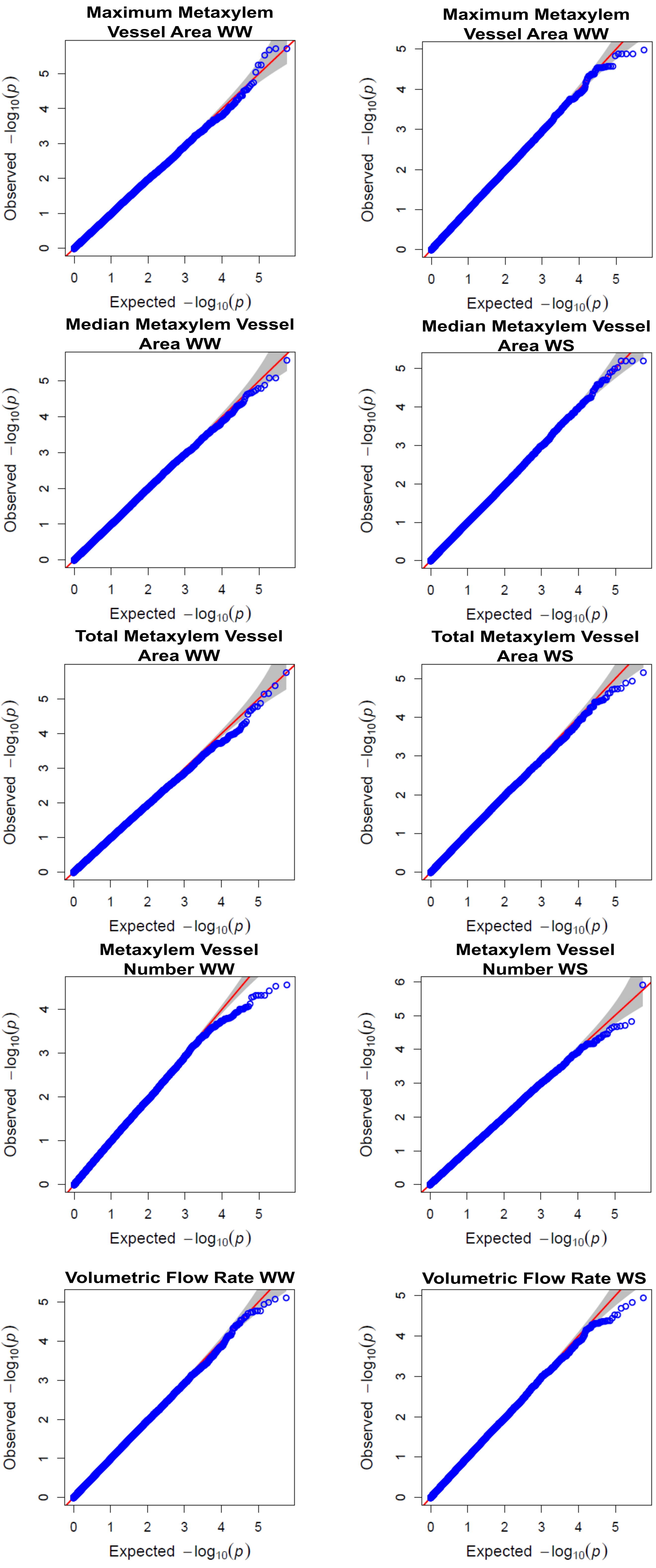


Supplemental Figure S2. Q-Q plots assessing the fitness of the model for GWAS of root metaxylem phenes under well-watered (WW) and water stress (WS) conditions.

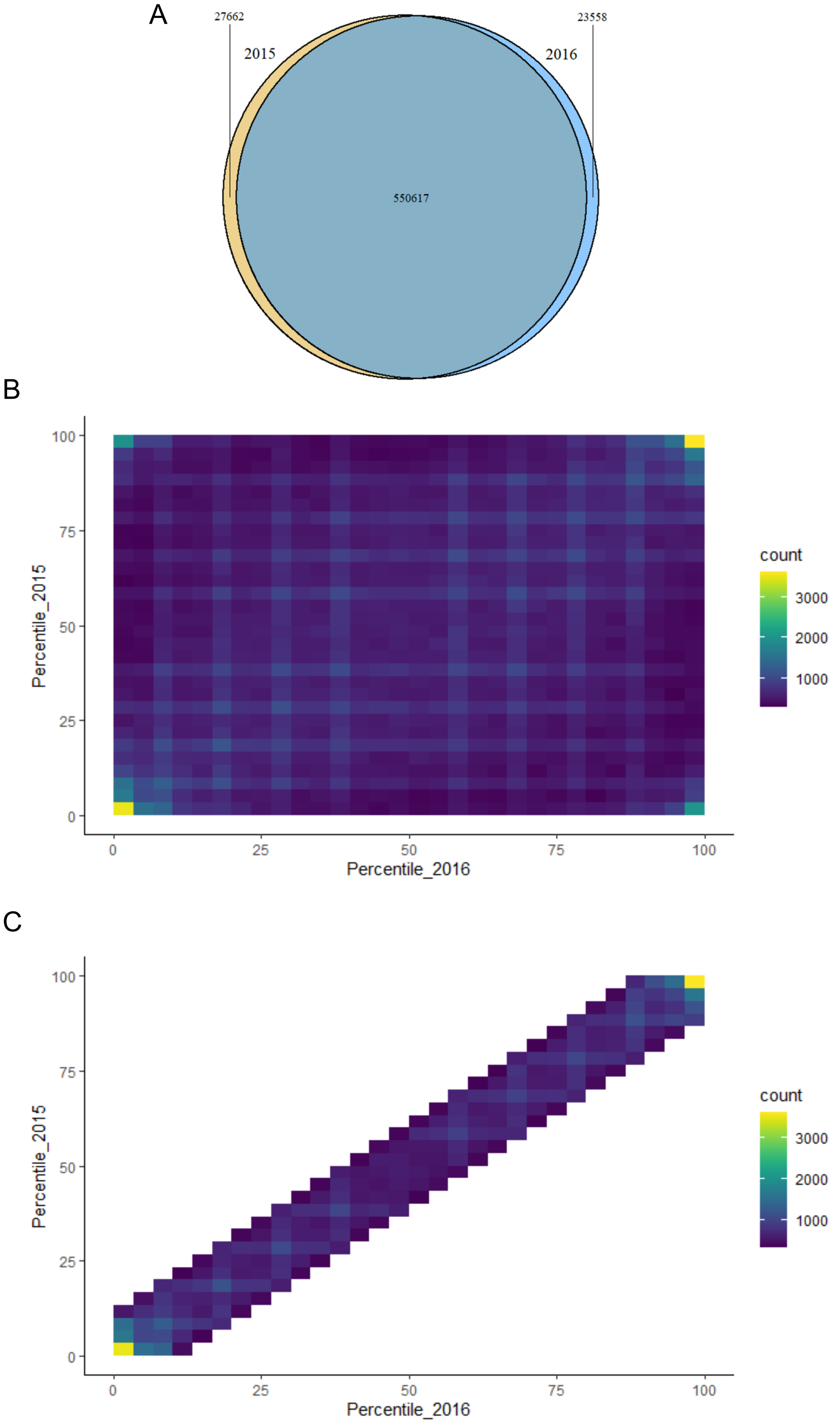


Supplemental Figure S3. General outline of the filtering steps used to identify SNPs with consistent allelic effects in separate GWAS conducted with data collected in 2015 and 2015. (A) A substantial proportion of the SNP panel was included in each GWAS. Binned scatterplots showing (B) the full SNP panel and (C) the filtered set containing SNPs of consistent minor allele effects. The gradation of the bin corresponds to the number of SNPs residing in the bin where densely populated bins are light blue while more sparsely populated bins are dark blue.

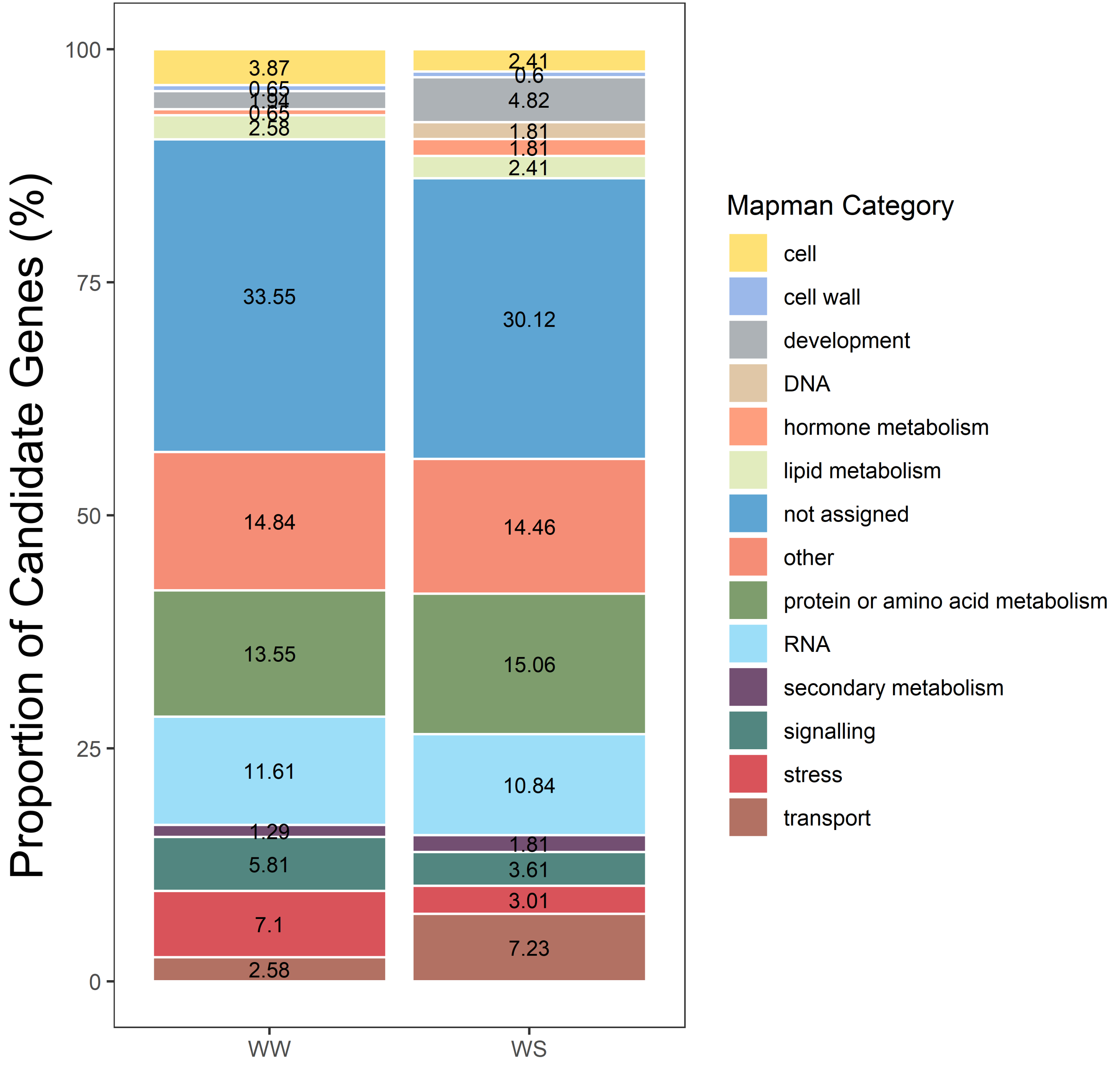


Supplemental Figure S4. Mapman ontogenic categories for annotated gene models associated with significant SNPs in well-watered and water stress conditions.

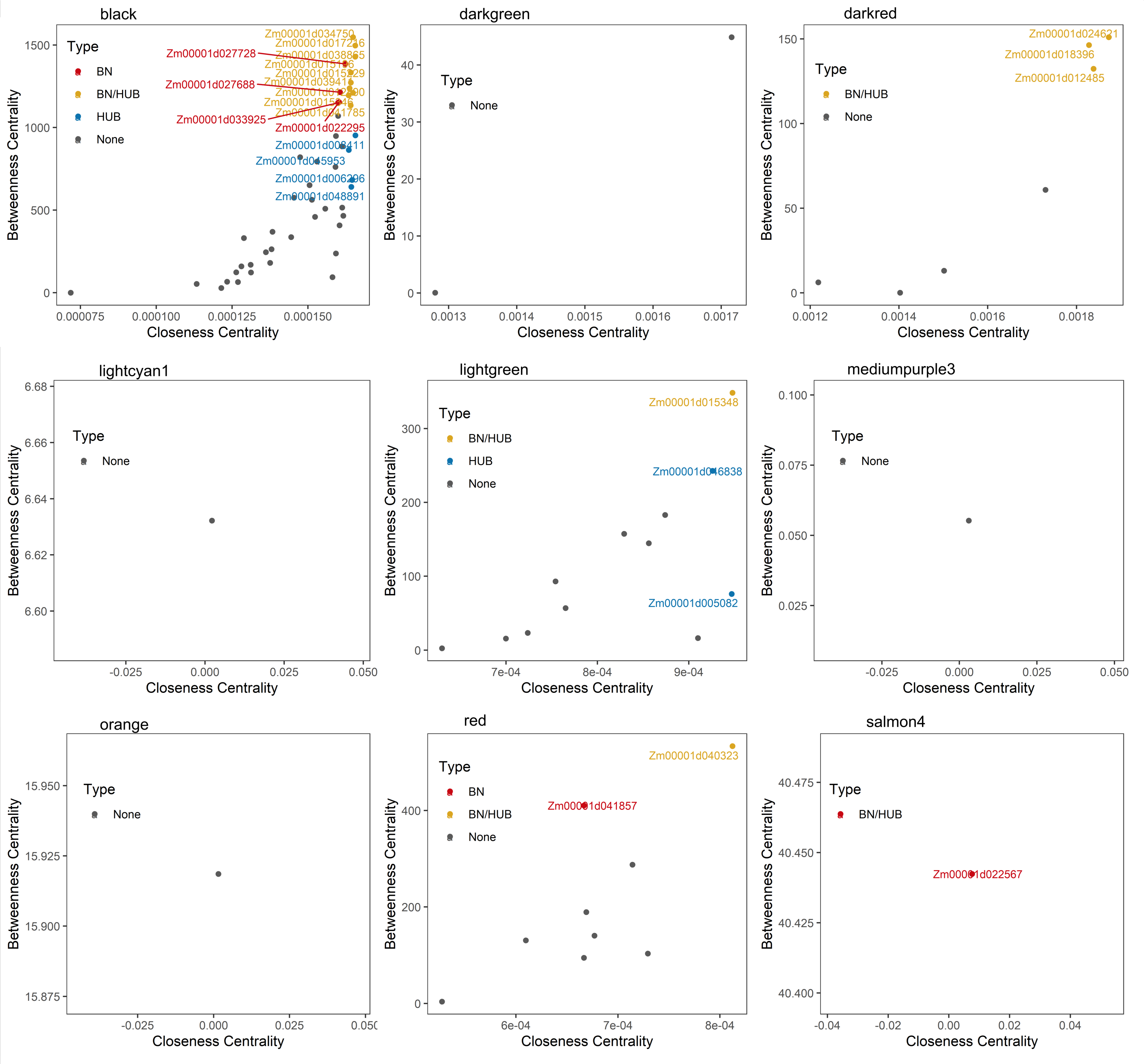


Supplemental Figure S5. Network topology metrics used to identify hub and bottleneck genes in the pool of candidate genes identified via GWAS. Scatterplots showing closeness centrality against betweenness centrality of each gene for the remaining gene co-expression modules most strongly associated with root tissues relevant to metaxylem development. Modules "navajowhite2" and "thistle1" did not contain any significant SNPs detected in the GWAS. Colors correspond to bottleneck (BN, red), bottleneck/hub (BN/HUB, yellow), hub (HUB, blue), and non-affiliated (None, grey) genes.

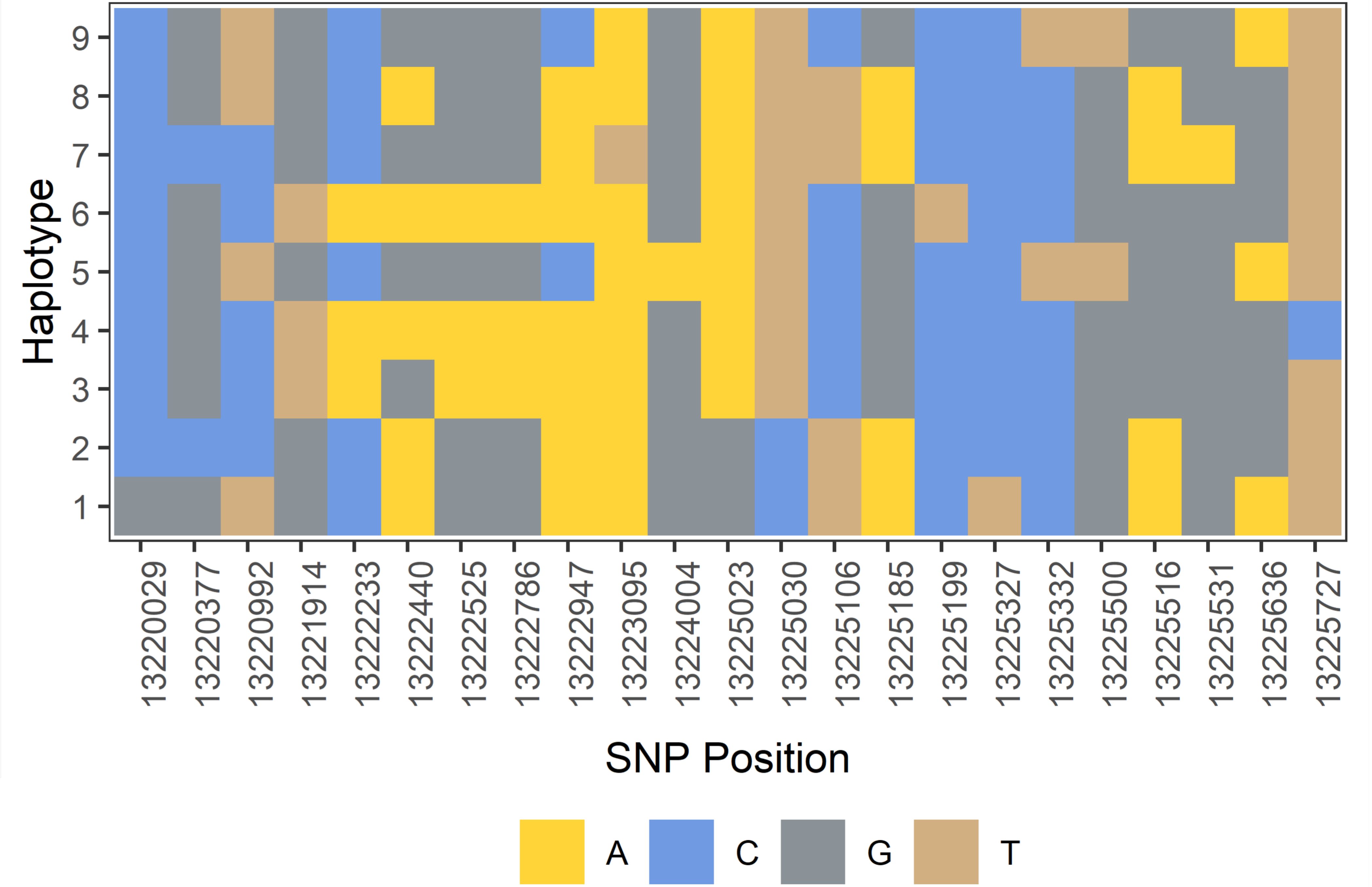


Supplemental Figure S6. Haplotypes of rice genes encoding a root-specific kinase 1 (Os10g0395000).
